# Tracking neural representations of attended and unattended features in multisensory working memory over time

**DOI:** 10.1101/2025.04.24.650418

**Authors:** Ceren Arslan, Daniel Schneider, Stephan Getzmann, Edmund Wascher, Laura-Isabelle Klatt

## Abstract

Working memory supports goal-directed behavior by maintaining task-relevant information. However, a growing number of studies shows that even task-irrelevant features are automatically encoded and can interfere with the recall of task-relevant information. While this phenomenon is well documented in unisensory contexts, it remains unclear whether and how task-irrelevant information persists in multisensory working memory. We examine the role of cross-modal binding by tracking the dynamic neural representation of audio-visual objects under varying selective attention conditions, using an EEG-based audio-visual delayed-match-to-sample task. Participants attended to either auditory or visual features (selective attention), or to both features (conjunction) of two sequential audio-visual items and subsequently compared those task-relevant features to an audio-visual probe. Further, to investigate the influence of bottom-up factors on cross-modal binding, we manipulated spatial congruency by presenting both features from either the same (i.e., in the center) or from disparate positions. Behaviorally, task-irrelevant features interfered with performance even under selective attention, consistent with automatic cross-modal binding and encoding into working memory. This interference was even stronger with spatially disparate presentation, suggesting that bottom-up attentional dynamics strengthen cross-modal bindings and their subsequent storage. Condition-level representational similarity analysis (RSA) showed that EEG activity patterns under selective attention more closely resembled those of conjunction trials than unisensory trials, indicating that task-irrelevant features were incorporated into multisensory object-level representations. This conjunction-similarity persisted in attend-visual trials, but declined over time in attend-auditory trials, reflecting partial filtering of task-irrelevant orientations. Crucially, activity patterns never shifted fully towards an auditory-only profile, indicating that irrelevant visual features were not fully excluded. In addition, feature-level RSA corroborated that both sound frequency-specific and orientation-specific information were present in EEG activity patterns, irrespective of the attended modality. Overall, these results demonstrate the persistence of task-irrelevant information in multisensory working memory and offer critical insights into how attentional processes shape its representational architecture.

## Introduction

Working memory is known to be fundamentally capacity limited (Cowan et al., 2008; Miller, 1956; Oberauer et al., 2016). Efficient use of these limited resources requires the selective prioritization of relevant information. Somewhat contrary to this assumption, a growing body of evidence demonstrates that task-irrelevant features are encoded into working memory in a largely automatic fashion (Bays et al., 2022; Marshall & Bays, 2013). However, the conditions under which these task-irrelevant features persist remain actively debated (e.g., Saltzmann et al., 2023; Shi et al., 2024; Woodman, 2021). Crucially, the existing literature has largely described this phenomenon within single modalities — primarily vision and audition — whereas it remains unclear whether automatic coding of task-irrelevant features into working memory takes place also when presented in a different modality. To address this gap, the present study investigates the fate of task-irrelevant features in multisensory working memory using both electroencephalography (EEG) and behavioral measures.

Behaviorally, the persistence of task-irrelevant features in working memory has often been indirectly assessed using irrelevant-change effects (Fischer et al., 2024; Gao et al., 2011; Shen et al., 2013). For instance, when comparing a probe stimulus, containing both task-relevant and task-irrelevant features, to working memory contents, performance on match trials decreases when a task-irrelevant probe feature changes, compared to when the probe presents a full match with the memory item. Analogously, rejecting a non-match-probe is less accurate or significantly slower when the task-relevant feature is paired with a matching task-irrelevant feature at test. Such “irrelevant change effects” have been shown in the visual modality for task-irrelevant changes of color, shape, and locations (Ecker et al., 2013; Gu et al., 2020; Logie et al., 2011), in audition, for task-irrelevant changes of frequency, location, and temporal amplitude modulation rate (Fischer et al., 2024; Joseph et al., 2015), and more recently in an audiovisual context, involving changes in the task-irrelevant modality (Arslan et al., 2025). Similarly, the event- and object-file literature – concerned with feature binding in light of object stability over time – reports analogous effects for auditory, visual, and multimodal bindings, which are typically referred to as ‘partial repetition costs’ (Noles et al., 2005.; Zmigrod et al., 2009; Zmigrod & Hommel, 2009, 2010). Together, these lines of research support the assumption that storage in working memory is – to some degree – object-based. Note that we have previously referred to such partial repetition costs (or irrelevant-change effects) as “probe congruency effects” (Arslan et al., 2025) and will continue to use this terminology here for consistency.

Neuroimaging studies provide additional, albeit mixed, evidence on the extent to which task-irrelevant features are represented in working memory. For example, Chen et al. (2021) were able to reconstruct task-irrelevant colors from multivariate electroencephalography (EEG) data, but these reconstructions were less reliable than those for task-relevant features.

Similarly, uncued colors were reconstructed from impulse-evoked EEG data (Kandemir et al., 2024), providing converging evidence that irrelevant visual features can persist, at least transiently or in activity-quiescent states. In contrast, several studies have reported decoding only of task-relevant visual features, without any detectable neural representation of task-irrelevant features (Bocincova & Johnson, 2019; Yu & Shim, 2017). Shedding light on the conditions under which task-irrelevant features may or may not persist, an fMRI study by Xu (2010) varied the number of shapes and colors in a memory display, when only memory for color was task-relevant. BOLD activity in an area associated with shape processing was sensitive to task-irrelevant variations of the number of shapes; but critically, this load-dependent modulation was only evident when encoding load for the task-relevant feature (i.e., color) was low. This suggests that the degree to which working memory storage is object-based, even when only certain features are task-relevant, may rely on available resources as well as task demands (see also Saltzmann et al., 2023). In sum, while the majority of studies favor the view that encoding is relatively inflexible, encompassing both task-relevant and task-irrelevant features, some voluntary control over unwanted information in working memory is retained (Bays et al., 2022; Saltzmann et al., 2023; Xu, 2010).

When objects are composed of features from multiple modalities, analogous principles of object-based encoding have been demonstrated. Critically, multisensory integration is guided by two key factors: spatial proximity and temporal synchrony (Stein & Meredith, 1993; Stein & Stanford, 2008). Consistent with an important role for temporal synchronization, sound localization judgments are biased toward the position of a visual stimulus when the auditory and visual stimuli are temporally aligned (Radeau & Bertelson, 1987; Wallace et al., 2004), and audiovisual signals are more likely integrated into a single percept when they occur within a certain temporal binding window (Colonius & Diederich, 2004; Lewald & Guski, 2003; van Wassenhove et al., 2007; Wallace et al., 2004). In addition, selective attention to one modality has been shown to spread to temporally synchronous but task-irrelevant features in another modality (Eimer et al., 2004; Molholm et al., 2007). Moreover, multisensory interactions appear to be maximal when cross-modal features are spatially congruent as compared to spatially incongruent (Gondan et al., 2005; Teder-Sälejärvi et al., 2005, but see also Spence, 2013). For instance, faciliatory effects on stimulus detection or localization for bimodal compared to unisensory stimuli (Delong & Noppeney, 2021; Frassinetti et al., 2002), as well as related enhancements of neuronal activity (Macaluso et al., 2000), have been shown to depend on the spatial congruency between inputs. Importantly, these cross-modal interactions are not merely additive, but stimulate unique percepts (Mcgurk & Macdonald, 1976; Shams et al., 2000; Thomas, 1941) and neural signatures that are qualitatively different from the mere summation of unisensory inputs (Giard & Peronnet, 1999; Senkowski et al., 2011; Stekelenburg & Vroomen, 2007; Talsma & Woldorff, 2005; Teder-Sälejärvi et al., 2002; reviewed by Calvert & Thesen, 2004). This warrants the assumption that perceptual cross-modal interactions may fundamentally influence what and how information is encoded and maintained in working memory.

In a recent study, adopting an audiovisual delayed-match-to-sample paradigm, we showed that concurrently presented auditory and visual features are encoded into working memory, even when participants are instructed to attend to only one modality (Arslan et al., 2025). Behaviorally, the probe congruency effect emerged irrespective of which features were attended (i.e., auditory, visual, or both), demonstrating the automatic encoding and persistence of all object features. Notably, these costs were largest in the conjunction condition (i.e., when both auditory and visual stimulus were task-relevant), but reduced in the selective attention condition (i.e., when only one modality was task-relevant), consistent with stronger cross-modal binding when both modalities were attended. However, on the neural level, we did not find a complementary signature of integrated object storage. Rather, modulations of oscillatory alpha power were consistent with domain-specific allocation of attentional resources during maintenance. In addition, traditional ERP correlates of unisensory working memory load did not provide evidence for the active maintenance of task-irrelevant features. While this may be consistent with quiescent states of working memory storage, alternative explanations remain plausible: For instance, cross-modal interactions during encoding may have qualitatively transformed neural signatures, rendering standard unisensory ERP correlates less sensitive. On a similar note, it could be that if working memory storage was primarily object-based, then load modulations on the feature level might be attenuated. Alternatively, it is also possible that task-irrelevant features were reactivated only at recall, explaining the behavioral probe congruency effect while accounting for the lack of a clear neural signature during maintenance. Hence, to more closely investigate how and under which conditions task-irrelevant features are stored in audiovisual working memory, the present study adopted a refined design and methodology.

Specifically, participants performed a delayed match-to-sample task while EEG was recorded. In three multisensory conditions, participants sequentially encoded two audio-visual objects, consisting of tones (varying in pitch) and teardrop stimuli (varying in orientation). They were instructed to selectively attend to either (a) the auditory features (attend-auditory, AA), (b) the visual features (attend-visual; AV), or (c) both (conjunction). At test, participants were instructed to compare an audiovisual probe with the memorized items, while limiting their judgement (match versus no-match) to only the task-relevant dimension. In addition, we systematically manipulated whether auditory and visual features were presented from the same central position (i.e., spatially compatible) or were spatially segregated. This allows us to investigate the interplay of bottom-up factors (spatial congruency) and top-down attentional demands in shaping multisensory working memory representations, as indexed by both behavioral and neural measures. Critically, we also introduced two unimodal baseline conditions in which participants encoded and retained two tones (auditory-only) or two orientations (visual-only). The latter are instrumental in applying representational similarity analysis (RSA) to assess the degree to which task-irrelevant features are filtered out or suppressed after initial automatic encoding. Specifically, we assessed whether and at what points in time, the neural activity patterns in attend-auditory and attend-visual conditions more closely resemble conjunction trials or their respective unisensory counterparts. If activity patterns in attend-auditory and attend-visual trials exhibit higher similarity to activity patterns in conjunction trials throughout the delay period, this would indicate integrated storage of both task-relevant and task-irrelevant features. In contrast, if the task-irrelevant features were filtered out after initial encoding, activity patterns should approach greater similarity to activity patterns in the respective unimodal condition. Moreover, we expected that the spatial presentation format would affect the extent to which task-irrelevant features are integrated and, thus, subsequently retained in working memory. That is, we hypothesized that cross-modal interactions should be maximal when auditory and visual features were spatially compatible, resulting in a larger probe congruency effect as well as greater similarity of neural activity patterns with conjunction trials compared to spatially segregated presentations.

## 2. Materials and Method

### 2.1 Ethics Statement

This project was approved by the Ethical Committee at the Leibniz Research Centre for Working Environment and Human Factors and was conducted in line with the Declaration of Helsinki. All participants provided written informed consent and were compensated for participation with either 12 euros per hour or course credits.

### 2.2 Participants

A total of 37 healthy, young adults participated in this study. One participant was excluded from the study due to performing the auditory tasks below the chance level, and one participant was discarded from the dataset due to excessive bridging between EEG electrodes. The final dataset consists of 35 participants (20 women, 15 men, three left-handed) with a mean age of 25.14 years (*SD* = 4.21, age range = 18 to 34).

A sensitivity analysis revealed that with a sample size of 35 participants in a 3 x 2 factorial repeated-measures design, 80% power, and an alpha-level of .05, we can detect effects of (𝜂^2^_*p*_ = .20 (i.e., *d* = 0.49).

Participants reported no history of psychopharmacological medication or neurological disorders. Further, we evaluated the hearing acuity of participants, presenting eleven pure-tone frequencies between 125 Hz and 8 kHz in a pure-tone audiometry (Oscilla USB 330; Inmedico, Lystrup, Denmark). All participants had normal hearing below or equal to 25 dB for frequencies between 125 Hz to 3 kHz, while seven participants showed slightly elevated hearing levels of 30 – 40 dB at frequencies ≥ 4 kHz.

All participants were tested for visual acuity using Landolt C optotypes at a 1.5 m distance. Calculated by a logarithmic averaging procedure (Bach & Kommerell, 1998), the average visual acuity was 1.41 (*SD* = 1.17, range = 0.94 to 1.88), indicating good vision.

### 2.3 Experimental setup and stimuli

The experiment was carried out in a sound-attenuated, dimly lit room (5.0 × 3.3 × 2.4 m³) with foam panels on the walls and ceiling and a woolen carpet on the floor. The stimulus presentation was implemented using E-Prime (v 3.0, Psychology Software Tools, Pittsburgh, PA). Furthermore, we used an AudioFile Stimulus Processor to control the synchronization between auditory stimuli and EEG triggers (Cambridge Research Systems, Rochester, UK).

Visual stimuli were presented on a 49’’ centrally aligned monitor with a 5120 by 1440-pixel resolution and a 100 Hz refresh rate (Samsung, Seoul, South Korea). Three full-range loudspeakers were mounted at azimuthal positions of -90°, 0°, +90° and at a height of ∼1.3 meters. Participants sat down in a comfortable chair 1.5 meters from the screen. To minimize head movements, participants rested their head on a chin rest throughout the recording.

As visual stimuli, a pool of eight teardrop-shaped orientations with angles between 22.5° and 337.5°, was generated using the MATLAB® (R2022a) function ‘polarplot’. A visual mask stimulus was created by overlaying all orientations (for more details about the experimental stimuli, see Arslan et al., 2025). Additionally, four colored fixation crosses were used, including white (RGB values 255, 255, 255), blue (RGB values 82, 171, 255), orange (RGB values 252, 146, 83), and a combination of the latter two colors, each of which subtended a visual angle of 0.38° both horizontally and vertically. For auditory stimuli, eight harmonic tones with base frequencies of 270, 381, 540, 763, 1080, 1527, 2159, and 3054 Hz were generated using Cool Edit 2000 1.1 (Syntrillium Software Corporation, Phoenix, USA). Each tone consisted of the following harmonics: first harmonic with the base frequency at 100% amplitude, the second at 40%, the third at 30%, the fourth at 20%, and the fifth at 10% amplitude. Tones had 10 ms fade-in and fade-out time windows with a sampling frequency of 44100 Hz. A normalized auditory mask stimulus was obtained by mixing all eight tones. Auditory stimuli were played at an average sound level of 76 dB (LAeq).

### 2.4 Procedure, task, and experimental design

Figure 1 illustrates the task’s sequence of events. A briefly flashing fixation cross (i.e., increasing in size from 0.38° to 0.76° for 100 ms) signalled the start of each trial and remained on screen for a variable duration between 800 and 1100 ms until the first memory item appeared. Then, participants were presented with either an auditory, a visual, or an audiovisual memory item for 500 ms. For audiovisual stimuli, the pairs of tones and orientations were randomly selected on a trial-by-trial basis. Audiovisual features were always temporally aligned, but could be either spatially compatible (50%, i.e., both presented from a central position) or spatially disparate (50%). In the latter, the visual orientation remained in the center and was paired with a sound from either the left or the right side. Analogously, in auditory-only trials the sounds were presented from a central (50%), the left (25%) or the right (25%) position, whereas in visual-only trials the orientations were always presented from the central position. All subsequent stimuli within a trial were presented in the same modality and at the same spatial locations as the initial memory item. To minimize the impact of sensory memory traces, each memory item was followed by either an auditory, a visual or an audio-visual mask for 200 ms after an inter-stimulus interval (ISI) of 200 ms. Subsequently, after a delay of 600 ms a second memory item and a second mask were presented in the same format as before (i.e., auditory, visual or audiovisual). Following another delay of 1000 ms, an auditory, a visual, or had 2000 ms to respond. Participants performed a delayed-match to sample task, indicating whether the task-relevant probe feature(s) matched their memory contents.

**Figure 1.**
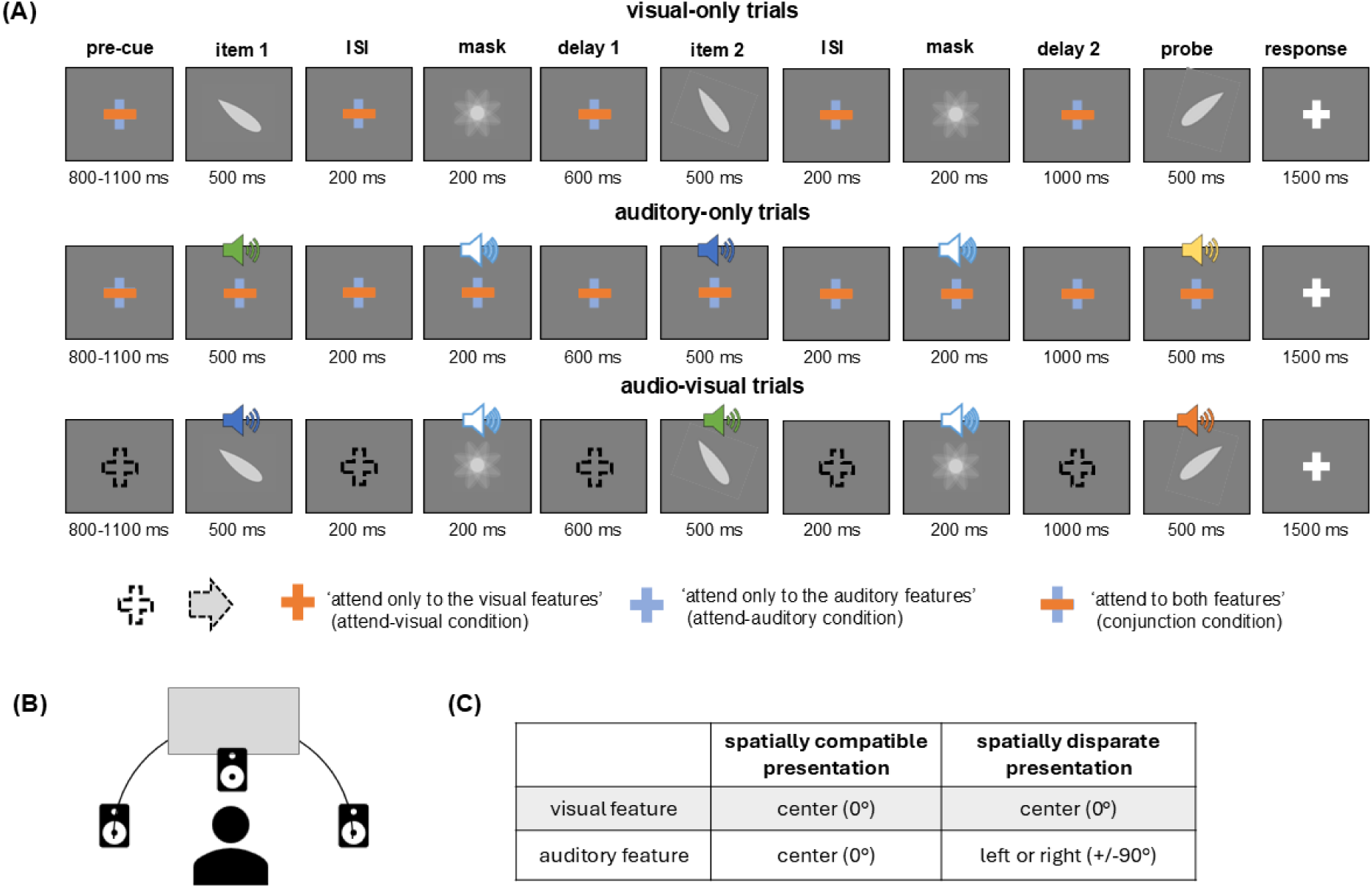
Schematic illustration of the trial sequence. (A) Throughout the experiment, participants were presented with two unisensory and three audio-visual trial-types. Each trial followed a delayed-match-to-sample format, in which participants encoded two sequentially presented memory items into working memory and subsequently judged whether a probe matched any of the items held in memory. In visual-only trials (V-only, top panel), participants were presented with teardrop shapes, varying in orientation. In auditory-only trials (A-only, middle panel), participants were presented with harmonic tones, varying in frequency. In audio-visual trials, both auditory and visual features were presented simultaneously and participants were instructed to attended to auditory features (blue fixation cross), visual features (orange fixation cross) or both features (bi-colored fixation cross, conjunction). The five conditions were grouped into two different types of blocks: In A-Blocks, conjunction, A-only, or V-only trials were presented in random order. Here, the fixation cross was always bi-colored, serving as a neutral cue. That is, participants were aware that all presented features, irrespective of their modality, will be task-relevant. In B-Blocks, attend-visual and attend-auditory trials were presented in random order. The color of the fixation cross reliably indicated the to-be-attended modality throughout the trial. (B) Experiment set up. A central loudspeaker was mounted below the screen at 0° azimuth. Two additional loudspeakers were mounted, one to the left (90°) and one to the right (+90°) of the participant.(C) By varying the stimulus positions we manipulated the spatially compatibility between auditory and visual features in audio-visual trial. The sounds were either spatially compatible with the centrally presented visual feature (50% of trials) or presented from a spatially disparate position (i.e., left or right, 25% each). Analogously, sounds were presented from the same three possible locations in auditory-only trials, while the visual features were always presented from a central position in visual-only trials.

The experiment consisted of five memory conditions (Figure 1A) that were grouped into two types of blocks: A-Blocks consisted of the two unisensory trial types (i.e., auditory-only [A-only], and visual-only [V-only] trials) as well as conjunction trials. In the latter, tones and orientations were presented concurrently, and participants were instructed to attend to both features. At the end of a trial, they compared these audio-visual objects to an audio-visual probe. In B-Blocks, participants were always presented with audiovisual information, but instructed to attend to either auditory (i.e., attend-auditory, AA trials) or visual (i.e., attend-visual, AV trials) features. At the beginning of each trial, the color of the fixation cross indicated the relevant modality and stayed on screen as a reminder: an orange fixation cross signaled attention to visual features, while a blue fixation cross directed them to focus on auditory features. At the end of a trial, they compared only the attended feature dimension of the probe to their memory contents.

Accuracy and speed were equally emphasized in all conditions. Responses were given by pressing one of the two horizontally arranged buttons on a response pad. The assignment of the left vs. right button with response options ‘yes’ vs. ‘no’ was counterbalanced across participants.

In all audiovisual conditions (attend-auditory, attend-visual, conjunction), probes could be either congruent or incongruent (Figure 2). *Congruent probes* consist of a “full match” or a “full mismatch”, i.e., the entire audiovisual pair matches one of the previously presented memory items or constitutes two new features from the stimulus set, respectively. *Incongruent probes* present a partial match in either modality (e.g., the frequency of the probe corresponds to the sound frequency of a previously presented memory item but is paired with a new orientation). Note that no re-combination probes were presented; instead, the non-matching probe feature was always randomly selected from the set of unimodal features that had not yet appeared in that trial. Critically, the proportion of congruent and incongruent trials per condition was adjusted to balance the number of “yes” and “no” responses. Consequently, the conjunction condition contained 50% of fully matching, congruent probes (i.e., the only trial type that required a yes-response), while the remaining 50% of trials were evenly distributed among the three other probe types (Figure 2B). In contrast, in attend-visual and attend-auditory conditions, each of the four probe types occurred in 25% of the trials (Figure 2A).

**Figure 2.**
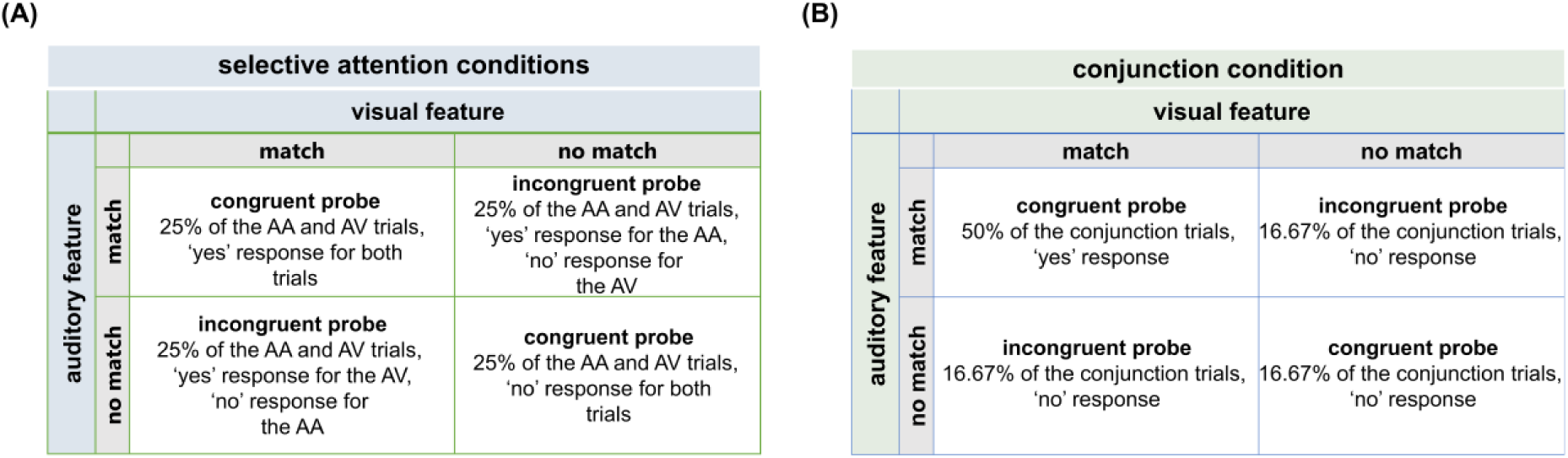
Probe types for attend-auditory (AA), attend-visual (AV), and conjunction trials. (A) In AA and AV, congruent and incongruent probes were presented equally often; (B) to equalize ‘yes’ and ‘no’ responses across probes in the conjunction condition, fully matching probes appeared in half of the conjunction trials, while the other three probes were equally presented across the remaining half of trials.

The grouping of memory conditions into two types of blocks was motivated by methodological considerations, allowing us to examine cross-modal interactions during encoding by means of an additive model framework. To this end, we additionally included a small proportion of aborted trials in both A- and B-Blocks, in which participants expected a regular trial to occur but in which no stimulus appears (Busse & Woldorff, 2003, Senkowski et al., 2011). Detailed explanations and corresponding results can be found in the supplementary material (section 1).

Overall, the experimental session was divided into four main blocks, including A-Blocks and B-Blocks in alternating order. The order of the blocks varied among participants, such that half of the participants were assigned the block order A-B-A-B, while the other half was assigned to the block order B-A-B-A. These main blocks were divided into 14 mini-blocks of approximately 6 minutes each: four mini-blocks in each A-Block and three in each B-Block. Across all blocks, each memory condition included 144 trials. A-Blocks consisted of a total of 432 main task trials and 192 aborted trials (i.e., no-stimulus events), while B-blocks included 288 main task trials and 126 no-stimulus events. Participants were given the opportunity to take self-paced breaks between mini-blocks. Before the start of the task, participants completed practice trials for each block type, lasting approximately three minutes. If the accuracy was below 50%, the practice was re-run. Overall, the experimental session (excluding the EEG preparation) lasted around two and a half hours.

### 2.5 EEG recording and preprocessing

EEG was recorded using a 64-channel Ag/AgCl electrode cap (BrainCap; Brainvision, Gilching, Germany), with electrode placement following the international 10-20 system. The signals were sampled at 1000 Hz using the NeurOne Tesla amplifier (Bittium Biosignals Ltd, Kuopio, Finland), and electrode impedances were maintained below 20 kΩ during cap preparation. AFz and FCz served as the ground and online reference electrode, respectively.

The data were pre-processed using MATLAB® (R2022a) and EEGLAB (2024.0; Delorme & Makeig, 2004). First, the continuous data were filtered with a 0.01 Hz high-pass (filter order: 330001, transition bandwidth: 0.01 Hz, −6 dB cutoff: 0.005 Hz) and a 40 Hz low-pass filter (filter order: 331, transition bandwidth: 10 Hz, −6 dB cutoff: 45 Hz). Channels showing normalized kurtosis exceeding five standard deviations from the mean were rejected using EEGLAB’s automated channel rejection tool (the pop_rejchan function). On average, 4.66 channels per participant were rejected (range: 1 to 8, *SD* = 1.70). The rejected channels were then interpolated using spherical splines based on neighboring channels. The data were re-referenced to the average of all channels.

Next, an independent component analysis (ICA) was conducted to facilitate artifact rejection. To optimize the data for ICA, in terms of processing speed and quality, the dataset underwent the following additional processing steps: First, the continuous data were down-sampled to 200 Hz for a faster ICA decomposition. In addition, data were filtered with a 1 Hz high-pass filter (Winkler et al., 2015). After segmenting the data into epochs between -1500 ms to 5500 ms relative to the first item’s onset, every other trial was selected to further reduce computation time (Getzmann et al., 2020; Klatt et al., 2020). Finally, an automatic trial-rejection method was employed to reject trials with large voltage fluctuations (> 1000 μV) and trials containing data values outside the range of three standard deviations (for details, see (documentation of *pop_autorej*).

The obtained ICA decomposition was then copied to the original continuous dataset (high-pass filtered at 0.01 Hz) with the primary sampling rate of 1 kHz and containing all trials. Again, the data were segmented into epochs spanning -1500 ms to 5500 ms relative to the onset of the first item and baseline-corrected using a pre-stimulus period from -100 to 0 ms. To identify and remove independent components (ICs) associated with non-brain activity, we applied the ICLabel plugin in EEGLAB (v1.4; Pion-Tonachini et al., 2019). ICs that reflected true brain activity with a probability of less than 30% were excluded. On average, 29.26 components (*SD* = 6.58) were removed per participant. Finally, any remaining trials showing large amplitude fluctuations (± 150 μV) were rejected.

After preprocessing, an average of 141.14 (*SD* = 3.84, 98%) auditory-only trials, 138.34 (*SD* = 8.27, 96%) attend-auditory trials, 140.46 (*SD* = 3.83, 97%) visual-only trials, 137.29 (*SD* = 9.06, 95%) attend-visual trials, and 141.31 (*SD* = 3.50, 98%) conjunction trials remained for further analysis.

### 2.6 Data analysis

Statistical tests were run using MATLAB® (R2022a) and JASP (version 0. 16.4.0). The significance of all tests was assessed at an alpha level of 0.05. We report partial eta squared (𝜂^2^_*p*_) and Cohen’s d (equation 10 in Lakens, 2013) as effect sizes for repeated measures ANOVA (rmANOVA) and t-tests, respectively. Bonferroni-Holm method was chosen to correct multiple comparisons, and corrected p-values are denoted as *pcorr*. Greenhouse-Geisser corrected *degrees of freedom* were reported if the assumption of sphericity was violated (Mauchley’s test *p* < .05). Representational similarity analyses (RSA) were run in Python (3.7).

#### 2.6.1 Behavioral analyses

To assess behavioral performance, accuracy (percentage of correct responses) and mean reaction times (RTs) were calculated per condition. Responses that occurred after 2000 ms following the onset of the probe were counted as incorrect responses (*M* = 0.001 trials).

First, to assess performance differences between all five memory conditions, separate one-way repeated measures analyses of variance (rmANOVA) were conducted for accuracy and RTs. These analyses disregarded the spatial compatibility factor, given that no such manipulation was present in the unisensory conditions. Second, to investigate the effects of the spatial manipulation and task conditions on overall performance, separate 3 x 2 rmANOVA were conducted for accuracy and RTs, including memory condition (attend-visual, attend-auditory, conjunction) and spatial manipulation (spatially compatible vs disparate) as within-subject factors. Third, to examine the probe congruency effect (i.e., the extent to which task-irrelevant features interfered at recall), accuracy and RT differences between congruent and (attend-visual, attend-auditory) and spatial compatibility (spatially compatible vs disparate). In addition, to verify whether the probe congruency effect was significantly different from zero within each condition (i.e., spatially compatible and disparate trials of attend-auditory and attend-visual conditions), individual one-sample t-tests were performed on the difference scores for both accuracy and RTs. Finally, exploratory paired sample t-tests were conducted to test whether the size of the probe congruency effect within each selective attention condition differed between spatially compatible and spatially disparate trials.

#### 2.6.2 Representational Similarity Analysis

Using representational similarity analysis (RSA; Kriegeskorte, 2008; Kriegeskorte & Kievit, 2013) we aimed to assess to what degree the task-irrelevant feature dimension is represented on a neural level when participants attend to only one modality. To this end, we conducted two sets of analyses (Figure 3A and B). First, we assessed the similarity of neural activity patterns between memory conditions. Here, the degree to which the attend-auditory or attend-visual trials exhibit stronger similarity to either their unisensory (auditory-only or visual-only) or multisensory (conjunction) counterparts allows us to determine whether the underlying representational structure more closely reflects individual features or integrated feature conjunctions. Second, an exploratory follow-up analysis aimed at assessing more explicitly to what degree feature-specific information concerning the task-irrelevant feature dimension is reflected in the neural activity patterns of attend-visual or attend-auditory trials. To this end, the similarity between trials of different feature values is compared.

**Figure 3.**
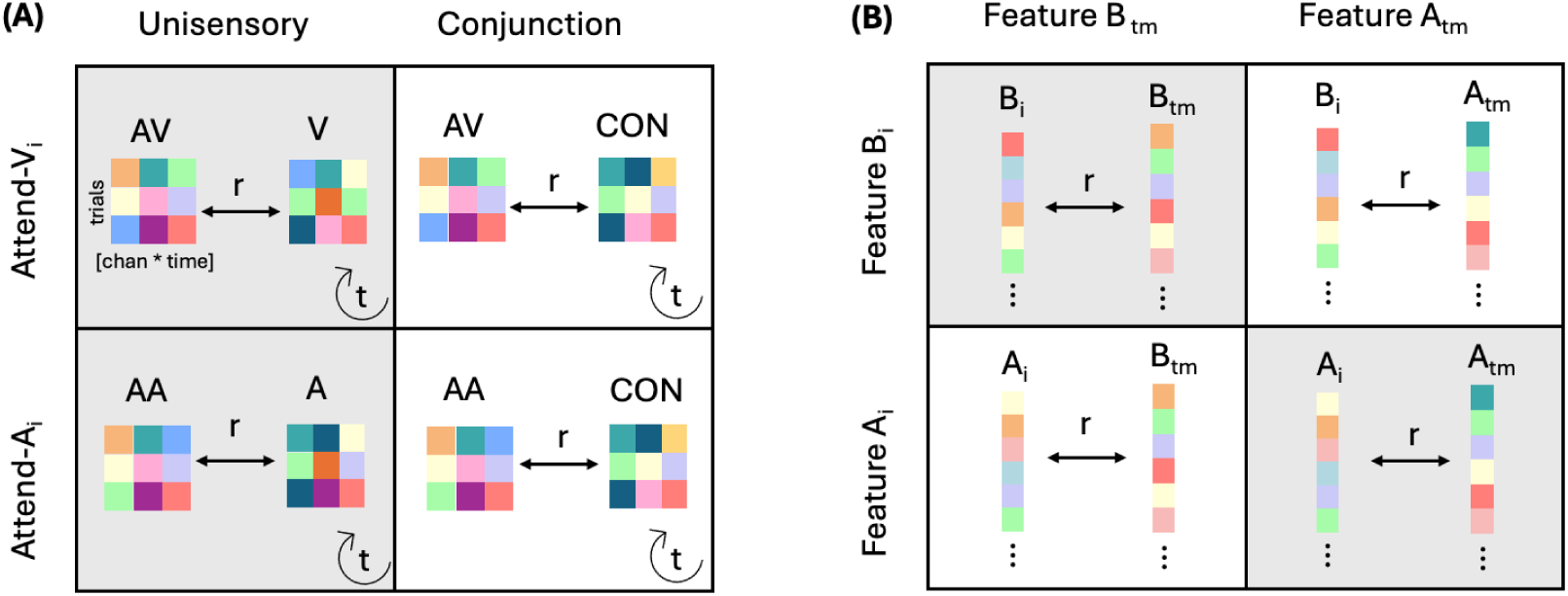
Schematic illustration of the conducted representational similarity analyses (RSA). (A) Condition-level RSA: The neural activity (channels x time bin) for each trial of one condition (e.g., attend-visual) is correlated with all other trials of another condition. The correlations were averaged across trials, resulting in one correlation estimate for each time bin. The procedure was repeated for all time points *t*, resulting in a time-series of similarity values. Specifically, we correlated trials from the attend-auditory and attend-visual condition with its unisensory control (grey) and the conjunction (white) condition. Each time bin contains 5 ms surrounding time point t. Accordingly, this process was iterated, using a sliding window approach. (B) Feature-level RSA: The flattened [channel x time] matrix for a given left-out trial *i* is correlated with the template [channel x time bin] matrix *tm* containing the average of all trials from the same category (grey) or a different category (white). That is, each colored square represents all time points in a defined window of interest (delay period; recall), appended in one row for all 64 channels. Features A and B represent either left-tilted vs. right-tilted orientations or high vs. low-pitched tones. The procedure is repeated, until all trials served as the left-out trial once.

##### 2.6.2.1 Representational similarity between task conditions

Representational similarity between conditions was assessed for the following condition pairs: (1a) attend-auditory and auditory-only trials (i.e., selective-unimodal-contrast), (1b) attend-auditory and conjunction trials (i.e., selective-conjunction-contrast), (2a) attend-visual and the visual-only trials (i.e., selective-unimodal-contrast), and (2b) attend-visual and conjunction trials (i.e., selective-conjunction-contrast). Subsequentially, the selective-unimodal and the selective-conjunction contrasts were compared (i.e., 1a vs. 1b and 2a vs. 2b) to assess whether the representations maintained in attend-auditory and attend-visual conditions more strongly resemble a “unisensory-like” structure or a “conjunction-like” structure. These steps were conducted separately for the spatially compatible and the spatially disparate condition. All analyses included correct trials only.

To obtain a time series of similarity scores, the following steps were performed for each condition pair: First, the trial x channel x time data matrix for each condition was obtained. A subset of trials was randomly selected if required to ensure an equal number of trials across conditions. Subsequently, using a sliding-window approach, the channel and time dimensions of the data were flattened for each condition within a time bin of 5 ms (n trials x [64 channels x 5 ms]). The neural activity for each trial of a condition was correlated with all trials of the contrasting condition using Spearman’s rank correlation coefficient (an approach adapted from Hucke et al., 2023). These correlations were averaged across trials, resulting in one correlation estimate (i.e., a single similarity score) for each time bin. This procedure was repeated for all time points within a trial (-1500 to 5500 ms relative to the first memory item).

The resulting similarity time series for each contrast (i.e., 1a, 1b, 2a, 2b) were compared using a non-parametric two-sided cluster-corrected sign-permutation test. For these tests, cluster_test() and cluster_test_helper() functions, provided by (Wolff et al., 2017) were used.

The cluster_test_helper() function produces a null distribution by randomly flipping the sign of each participant’s input data. This was repeated 10,000 times. The null distribution was input to the cluster_test() function, identifying clusters in the original data greater than the null distribution. The cluster-forming threshold and the cluster significance threshold were *p* < .05. Additionally, the difference between the similarities of selective unimodal and selective conjunction contrasts (i.e., 1a vs. 1b and 2a vs. 2b) was compared between spatially compatible and disparate conditions. Furthermore, the difference between the similarities of selective unimodal and selective conjunction contrasts in the visual modality (i.e., 1a – 1b) was tested against the similarity differences in the auditory modality (i.e., 2a – 2b), separately for spatially compatible and disparate trials. The results were tested using two-sided cluster-corrected sign-permutation tests.

##### 2.6.2.2 Representational similarity between features

To test if task-irrelevant features were retained in working memory, we assessed the similarity of neural activity patterns associated with individual feature values in attend-visual and attend-auditory trials. If orientations were maintained in working memory despite the instruction to focus on the sounds, then activity patterns for right-tilted orientations should be more similar to each other than to left-tilted orientations (or vice versa). Similarly, if the tones were represented in working memory in attend-visual trials, we expected activity patterns for trials with low-frequency tones to be more similar to each other than to high-frequency tones (or vice versa). Thus, all trials were labelled according to their task-irrelevant feature dimension for the first and the second memory item. Orientations were classified as right-tilted (i.e., 22.5°, 67.5°, 112.5°, 157.5°) vs. left-tilted trials (i.e., 202.5°, 247.5°, 292.5°, and 337.5°) and tones were classified as low (i.e., 270, 381, 540, 763 Hz) vs. high (i.e., 1080, 1527, 2159, and 3054 Hz) in frequency. We compared neural activity patterns in two time-windows of interest that were derived in a data-driven manner based on the first set of analyses (i.e., representational similarity between conditions), that is the delay period (2.4 to 3.4 s relative to the first memory item’s onset) and the probe-response period (3.4 to 4.2 s relative to the first memory item’s onset). Within those time windows, the data was flattened to obtain a trial x (channel * time point) matrix. Based on this data matrix, the representational strength of task-irrelevant auditory and visual features was quantified using a template-based leave-one-out approach (Günseli & Aly, 2020; Mumford et al., 2014): First, averaging all but one trial in a given category (e.g., left-tilted trials), we created a template activity pattern for that category (i.e., “same template”). Analogously, all trials belonging to the other category (e.g., right-tilted trials) were averaged, constituting the “different template”. The left-out trial (1x [channel*time point]) was then correlated with both the “same template” and the “different template”, using Spearman’s rank correlation coefficient. This procedure was repeated until all trials served as a left-out-trial

once and for both feature values (e.g., right-tiled and left-tilted trials). The resulting correlations were Fisher-transformed to ensure normality. Finally, correlations were averaged across iterations of left-out-trials and feature values (e.g., left- and right-tilted orientations), resulting in one “same” and one “different” similarity score for per participant. Difference scores (i.e., same-different correlations) were then compared to zero using one-sample t-tests, for both the tones and orientations. If the difference score is larger than zero, the neural activity pattern contains orientation- or tone-specific information. Based on the observation that activity patterns in attend-visual trials retained higher similarity to conjunction trials (compared to visual-only trials) throughout maintenance and recall, whereas this was not the case for attend-auditory trials, we further assessed if tones were more strongly represented in attend-visual trials than orientations in the attend-auditory trials. To this end, we conducted an additional paired-sample t-test contrasting the difference scores for attend-visual tones and attend-auditory orientations. A final set of paired-sample t-tests aimed to assess whether task-relevant features were more strongly represented than task-irrelevant features within each selective attention condition. All the tests described in this section were done separately for the first and second memory item during both the delay and probe response periods.

## 3. Results

### 3.1 Behavioral results

#### 3.1.1. Memory conditions

Figure 4 shows the overall behavioral performance observed for each memory condition, collapsed across spatially compatible and disparate trials. Our results showed no differences between conditions, neither in accuracies (*F*(3.31, 112.54) = 1.58, *p* = 0.19, 𝜂^2^_*p*_ = 0.05) nor in RTs (*F*(3.24, 110) = 2.22, *p* = 0.09, 𝜂^2^_*p*_ = 0.06).

**Figure 4.**
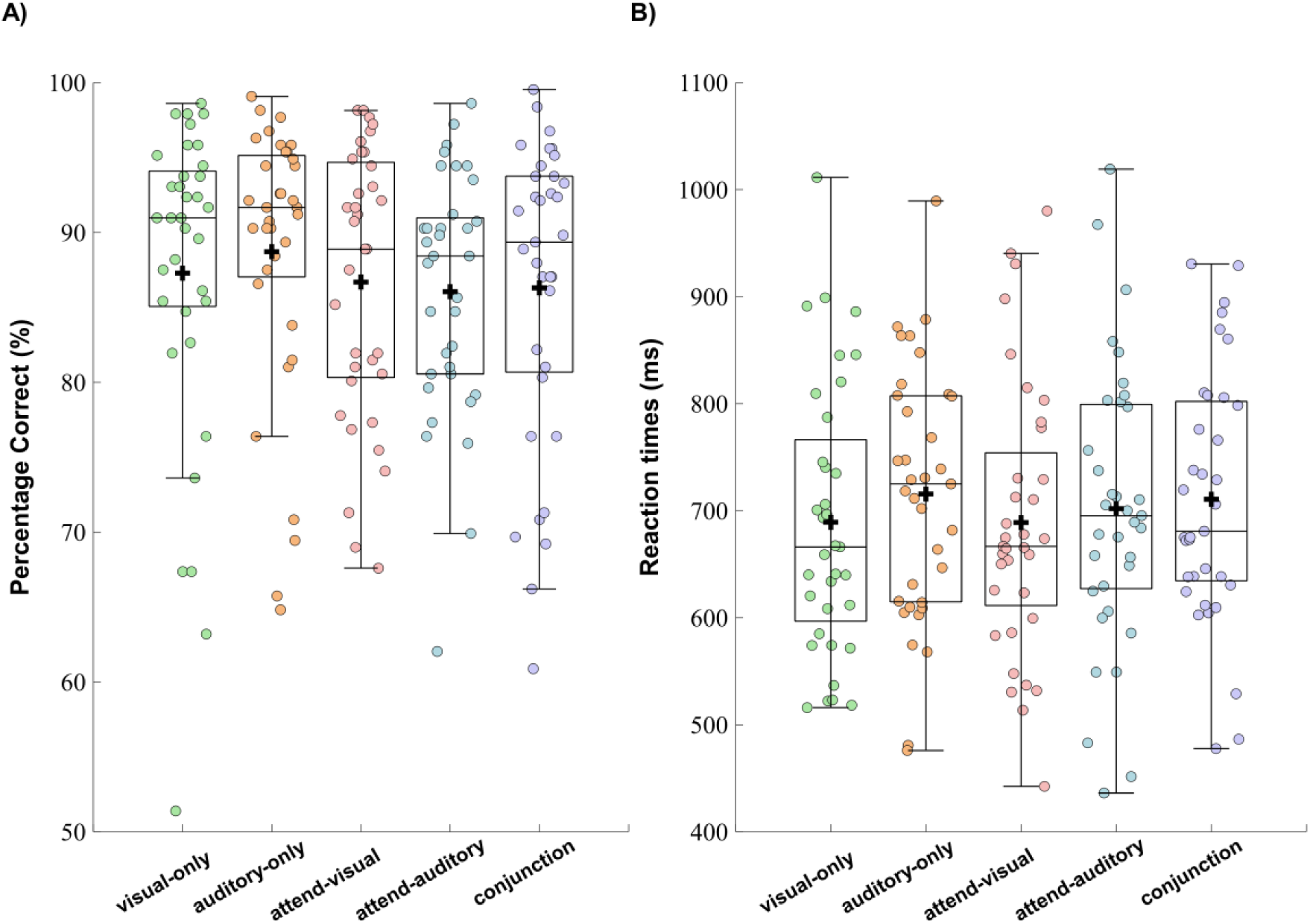
Behavioral performance across memory conditions collapsed across spatially compatible and disparate trials. (A) The proportion of correct responses; (B) RTs for the conditions. Boxplots show the +/- 1.5 interquartile range and the median. The dots illustrate individual participant averages per condition. A black cross illustrates the condition mean.

#### 3.1.2. The effect of spatial compatibility

Figure 5 illustrates behavioral performance for the three audiovisual memory conditions (attend-auditory, attend-visual, conjunction), separately for spatially compatible and disparate trials. For accuracy, the analysis revealed no main effect of memory condition (*F*(1.92, 65.32) = 0.46, *p* = .62, 𝜂^2^_*p*_ = 0.01) or spatial manipulation (*F*(1, 34) = 0.65, *p* = .43, 𝜂^2^_*p*_ = 0.02). However, a significant interaction between condition and spatial manipulation emerged (*F*(1.7, significantly higher accuracy than disparate trials in the attend-visual condition (*t*(34) = 3.04, *pcorr* = .02, *d* = 0.51). Thus, task-irrelevant lateralized tones interfered with task performance more strongly than centrally presented tones. In contrast, accuracy did not differ between spatially compatible and spatially incompatible trials in either the attend-auditory (*t*(34) = -1.22, *pcorr* = .46, *d* = -0.21) or the conjunction condition (*t*(34) = 0.50, p*corr* = .62, d = 0.08) (Figure 5A).

**Figure 5.**
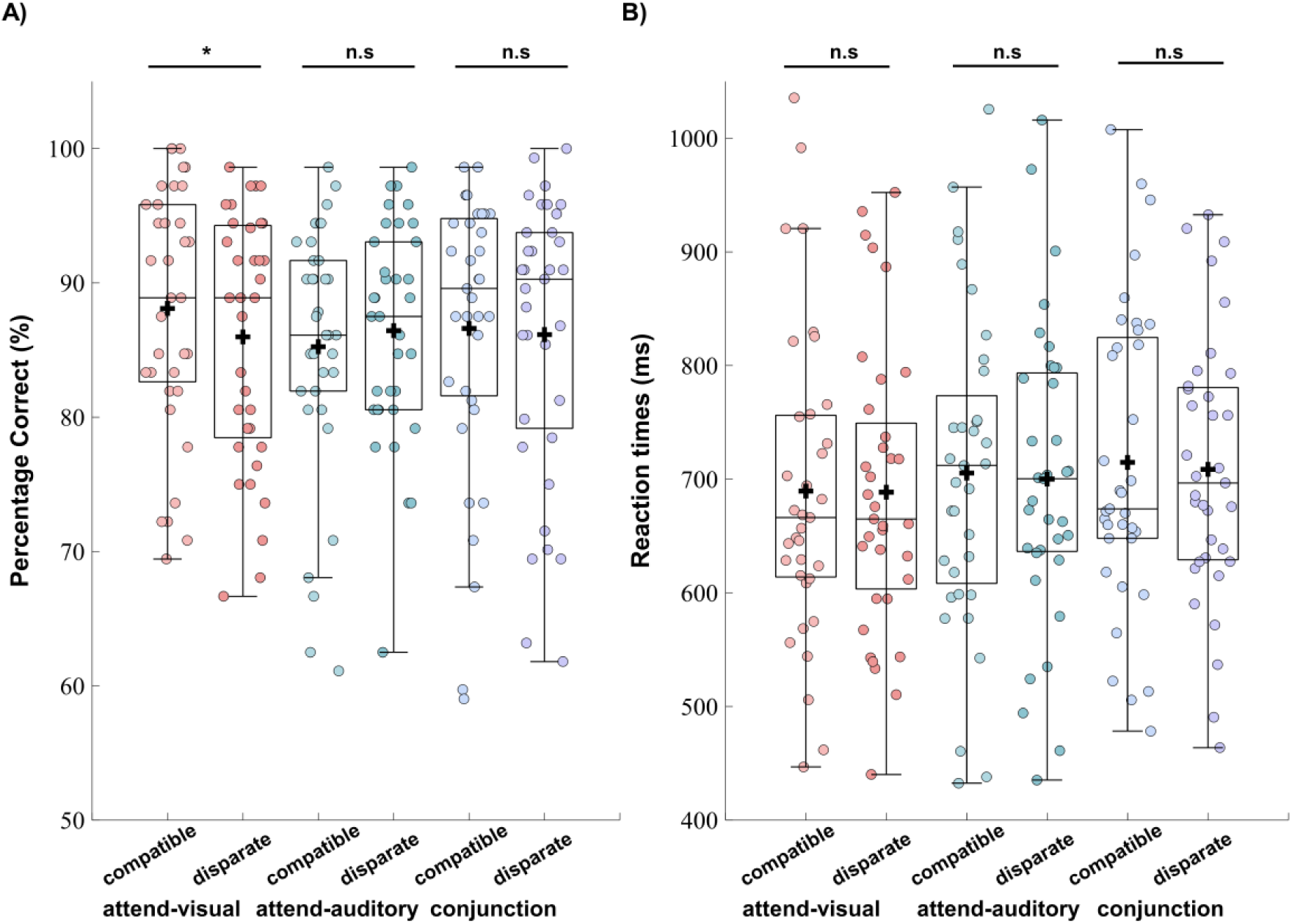
Behavioral performance across memory conditions for spatially compatible and disparate trials. (A) The proportion of correct responses; (B) RTs for the conditions. Boxplots show the +/- 1.5 interquartile range and the median. The dots illustrate individual participant averages per condition. A black cross illustrates the condition mean. \**p* <0.05, n.s=not significant.

For RTs, our analyses did not reveal a main effect of condition (*F*(1.7, 57.5) = 2.08, *p* = 0.14, 𝜂^2^_*p*_ = 0.06) or spatial manipulation (*F*(1, 34) = 0.45, *p* = 0.51, 𝜂^2^_*p*_ = 0.01), nor an interaction between these two factors (*F*(1.71, 58.03) = 0.08, *p* = 0.9, 𝜂^2^_*p*_ = 0.002) (Figure 5B).

#### 3.1.3. Probe congruency effect

Figure 6 illustrates the probe congruency effect in accuracy and response times separately for attend-visual and attend-auditory trials as well as spatially compatible and spatially disparate trials. The analysis revealed a main effect of spatial compatibility on accuracy (*F*(1, 34) = 6.52, *p* = .02, 𝜂^2^_*p*_ = 0.16), indicating a greater accuracy difference between congruent vs incongruent probes in the spatially disparate than compatible trials (Figure 6A). Yet, there was no main effect of condition (*F*(1, 34) = 0.38, *p* = .54, 𝜂^2^_*p*_ = 0.01) or an interaction between two factors (*F*(1, 34) = 1.02, *p* = .32, 𝜂^2^_*p*_ = 0.03).

**Figure 6.**
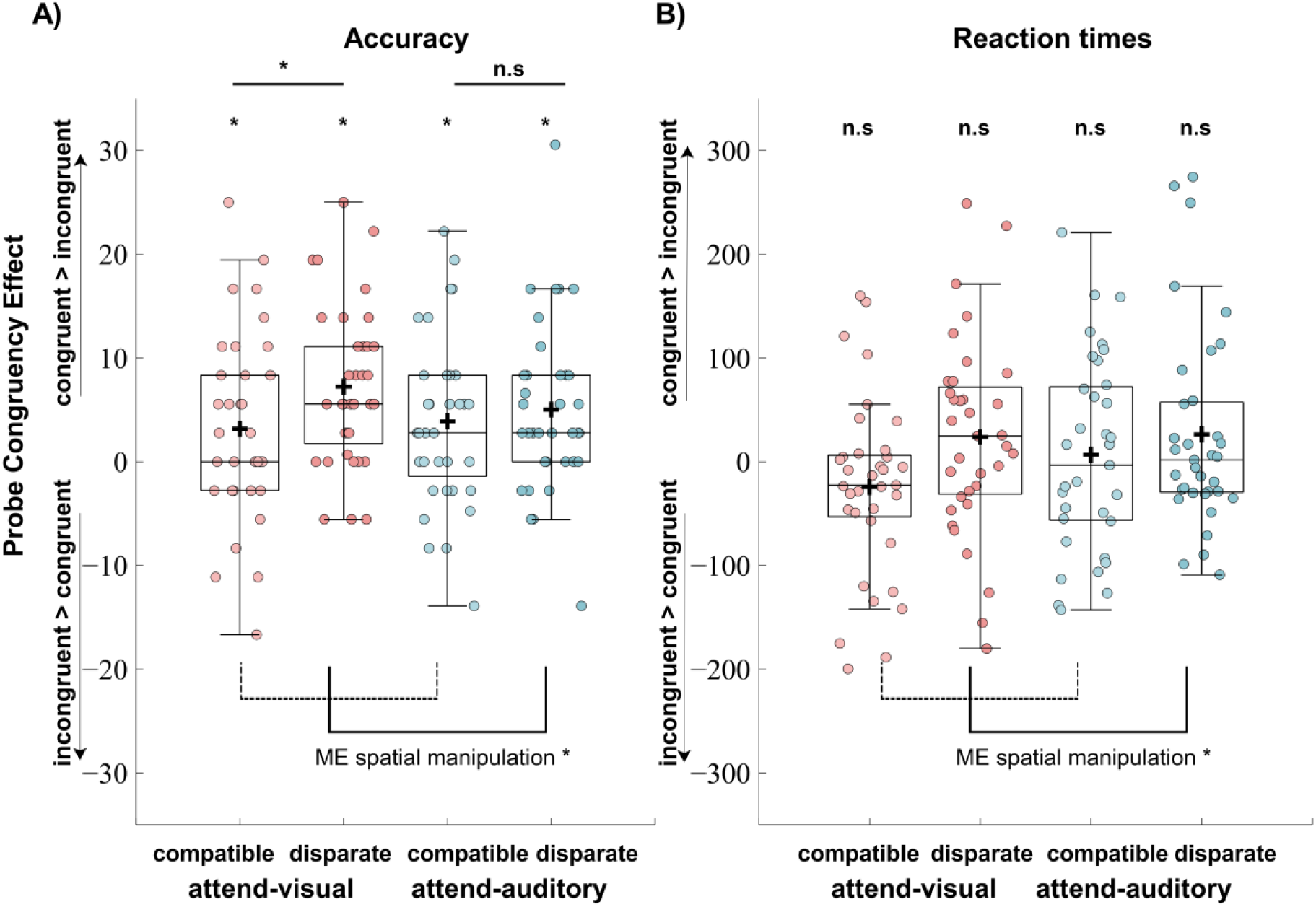
Probe congruency effects for each experimental condition. The probe congruency effect is quantified as the difference in proportion correct (A) and response times (B) between congruent and incongruent probe trials. Boxplots show the +/- 1.5 interquartile range and the median. The dots illustrate individual participant averages per condition. A black cross illustrates the condition mean. \**p* <0.05, n.s = not significant.

To verify the presence of a probe congruency effect for each condition, individual one-sample t-tests contrasted the difference scores (congruent – incongruent) against zero. The results showed higher accuracy for congruent probes compared to incongruent probes in attend-auditory trials, irrespective of whether audio-visual feature were spatially disparate (M = 5.03, SD = 8.22, *t*(34) = 3.62, *pcorr* = .004, *d* = 0.62) or spatially compatible (M= 3.91, SD = 8.15, *t*(34) = 2.84, *pcorr* = .02, *d* = 0.48). In attend-visual trials, a significant probe congruency effect also emerged in both spatially-disparate trials (M = 7.24, SD = 7.67, *t*(34) = 5.59, *pcorr* = .003, *d* = 0.94), as well as compatible trials (M = 3.18, SD = 9.04, *t*(34) = 2.1, *pcorr* = .045, *d* = 0.35), albeit only approaching conventional statistical thresholds in the latter.

The descriptively stronger effect of the spatial manipulation in attend-visual trials motivated exploratory comparisons of spatially compatible vs spatially disparate trials within each selective attention condition. Results corroborated that the probe congruency effect was significantly larger in spatially disparate than compatible trials of the attend-visual condition (*t*(34) = -2.41, *pcorr* = 0.04, d = -0.41). In contrast, there was no such difference in the attend-auditory condition (*t*(34) = -0.60, *pcorr* = 0.56, *d* = -0.10).

RT analyses revealed a main effect of spatial compatibility (*F*(1, 34) = 5.79, *p* = .02, 𝜂^2^_*p*_ = 0.15), indicating a greater RT difference between congruent vs incongruent probes in spatially disparate than compatible trials (Figure 6B). Yet, there was no main effect of condition (*F*(1, 34) = 1.26, *p* = .27, 𝜂^2^_*p*_ = 0.04) or an interaction between the two factors (*F*(1, 34) = 0.81, *p* = .37, 𝜂^2^_*p*_ = 0.02).

Unlike the accuracy results, the probe congruency effect within individual conditions was not established for RTs. There were no differences between congruent and incongruent probes in spatially compatible (*t*(34) = -1.67, *pcorr* = 0.4, d = -0.28) and disparate trials (*t*(34) = 1.49, *pcorr* = 0.3, d = 0.25) of attend-visual condition; and in compatible (*t*(34) = 0.42, *pcorr* = 0.67, d = 0.07) and disparate trials (*t*(34) = 1.61, *pcorr* = 0.36, d = 0.27) of attend-auditory condition.

Overall, in contrast to what we expected, we found a greater probe congruency effect for spatially disparate compared to spatially compatible stimulus presentation. Exploratory follow-up tests revealed that this was mostly apparent in the attend-visual condition, while the probe congruency effect was not modulated by the spatial manipulation in the attend-auditory condition. These findings suggest that, task-irrelevant, lateralized tones in attend-visual trials presented orientations in attend-auditory trials seems to be unaffected by the position of the tones.

### 3.2 Representational similarity between conditions

Figure 7 shows the similarity scores between memory conditions, separately for spatially congruent and disparate trials. For spatially compatible trials, RSA revealed significantly higher pattern similarity of attend-auditory trials with the conjunction than with the auditory-only condition. Significant clusters (*p* < .05) emerged during encoding as well as parts of the delay interval (i.e., 80 to 1210 ms, between 1890 to 2070 ms, and between 2460 to 2690 ms (Figure 7A). For spatially disparate feature presentation, a similar pattern emerged with significant clusters ranging from 70 to 1230 ms, 1760 to 2270 ms, and 2460 to 2770 ms (*p* < .05) (Figure 7B). Yet, the similarity differences between the selective unimodal and selective conjunction contrasts were not significantly different between spatially compatible and disparate trials (all *p* > .05). These results indicate that task-irrelevant orientations in the attend-auditory condition are represented in working memory until the second half of the delay period. In later parts of the trial, neural activity patterns were equally similar to the auditory-only and the conjunction condition, suggesting that task-irrelevant tones were partially filtered out but still weakly represented.

**Figure 7.**
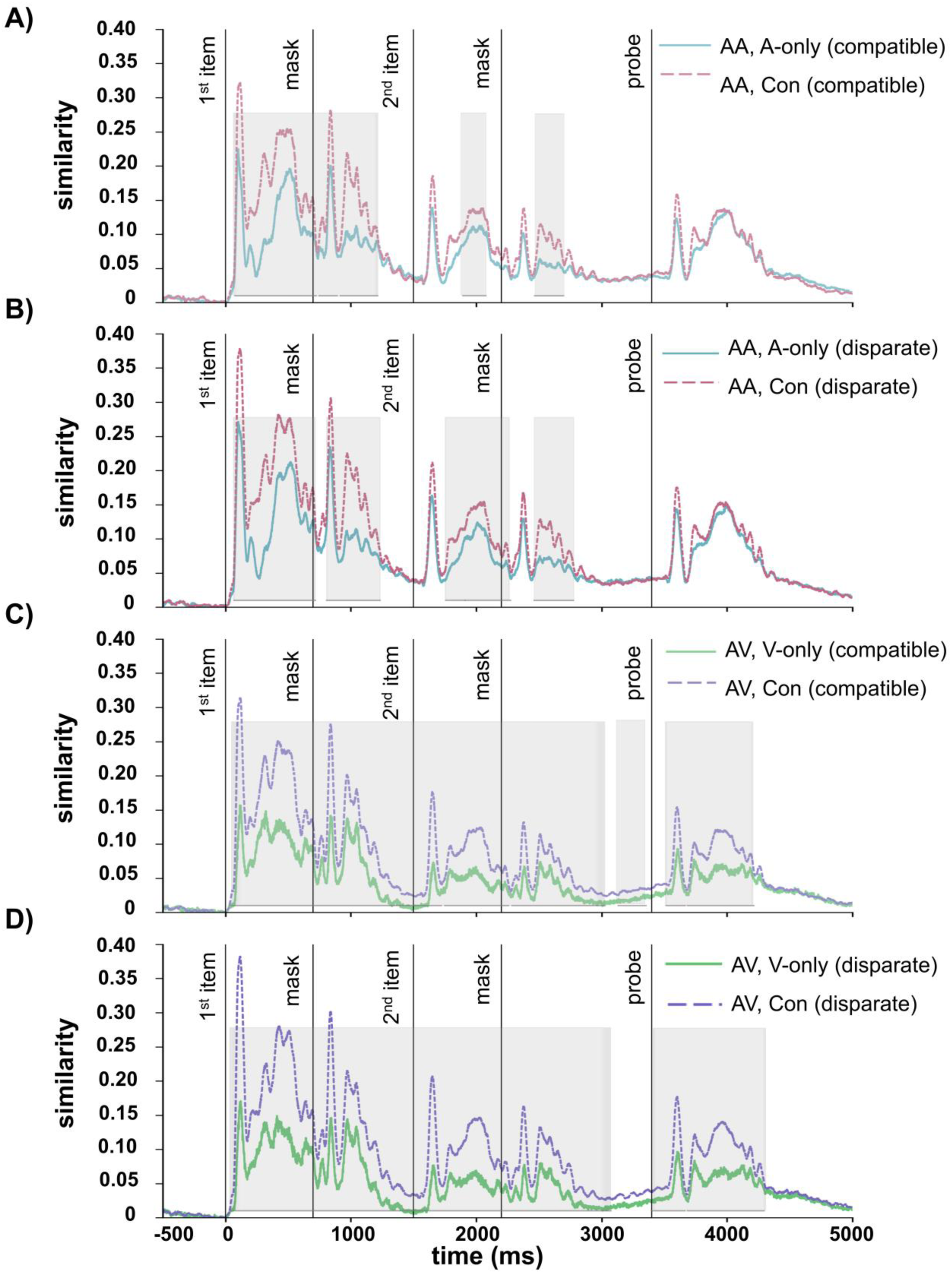
Similarities of neural activity patterns between task conditions for spatially compatible and disparate trials. Panels A and B show Spearman correlations between attend-auditory (AA) and auditory-only (A-only) trials (blue solid line) and between attend-auditory (AA) and conjunction (Con) trials (pink dashed line) for spatially compatible and for spatially disparate conditions, respectively. Panels C and D show between attend-visual (AV) and visual-only (V-only) trials (green solid line) and between attend-visual (AV) and conjunction (Con) trials (purple dashed line) for (C) spatially compatible and (D) for spatially disparate trials. Shaded areas show the time periods in which the pairwise comparisons significantly differed.

For spatially compatible attend-visual trials, RSA revealed higher similarity to the conjunction condition than to the visual-only condition. The effect was sustained throughout the majority of the trial with significant clusters ranging from 30 to 3350 ms and between 3520 to 4210 ms (*p* < .05) (Figure 7C). A compatible pattern was obtained for spatially disparate trials with significant clusters ranging from 40 to 3070 ms and from 3400 to 4300 ms (*p* < .05) (Figure 7D). Additionally, our results showed that the similarity differences between selective unimodal and selective conjunction contrasts were significantly higher in spatially disparate than compatible trials. The effect was sustained between 75 ms to 160 ms and 790 to 890 ms, 1590 to 1640 ms, 1970 to 2130 ms, 3560 to 3590 ms, and between 3945 to 3970 ms (*p* < .05). In sum, these results indicate that, unlike task-irrelevant orientations in attend-auditory trials, task-irrelevant tones in attend-visual trials appear to be maintained throughout the delay period and subsequent processing stages, resulting in persistently higher similarity with the conjunction condition. This effect was more pronounced when audiovisual features were spatially disparate than spatially compatible.

Lastly, corroborating those differential dynamics in the two selective attention conditions, the difference between the selective unimodal (AV and V-only) and selective conjunction contrasts (AV and conjunction) in the visual modality (Figure 7C) was significantly greater than the difference between those two contrasts in the auditory modality (i.e., AA and A-only vs. AA and Conjunction; Figure 7A) in spatially compatible trials. This difference was sustained with significant clusters between 70 to 110 ms, 460 to 580 ms, 800 to 900 ms, 1150 to 1670 ms, 1900 to 2110 ms, and between 3900 to 4120 ms (*p* < .05). Similarly, there was a greater difference between the similarities of selective unimodal and selective conjunction contrasts in the visual modality (Figure 6D) than in the auditory modality (Figure 6B) in spatially disparate trials, with sustained clusters between 70 to 155 ms, 250 to 310 ms, 400 to 615 ms, 790 to 890 ms, 1140 to 1700 ms, 1890 to 2140 ms, 2330 to 2410 ms, 2850 to 4130 ms (*p* < .05).

### 3.3. Representational similarity *between features*

Differences in the similarity of neural activity patterns to a same-feature-template compared to a different-feature-template served as an indicator of feature-specific representations in working memory (see Methods). One-sample t-tests showed that the difference scores (same vs. different correlations) for orientations of the first memory item were significantly different from zero in attend-auditory trials (delay: *t*(34) = 10.95, *pcorr* = 0.004, *d* = 1.85; probe-response: *t*(34) = 10, *pcorr* = 0.004, *d* = 1.69) and attend-visual trials (delay: *t*(34) = 3.69, *pcorr* = 0.003, *d* = 0.62; probe-response: *t*(34) = 7.85, *pcorr* = 0.003, *d* = 1.33), providing evidence for orientation-specific information in the neural activity patterns (Figure 8). The same was true for difference scores obtained for orientations of the second memory item in attend-auditory trials (delay: *t*(34) = 11.35, *pcorr* = 0.002, *d* = 1.92; probe-response: *t*(34) = 11.83, *pcorr* = 0.002, *d* = 1.98) and attend-visual trials (delay: *t*(34) = 11.26, *pcorr* = 0.001, *d* = 1.90; probe-response: *t*(34) = 10.24, *pcorr* = 0.001, *d* = 1.73).

**Figure 8.**
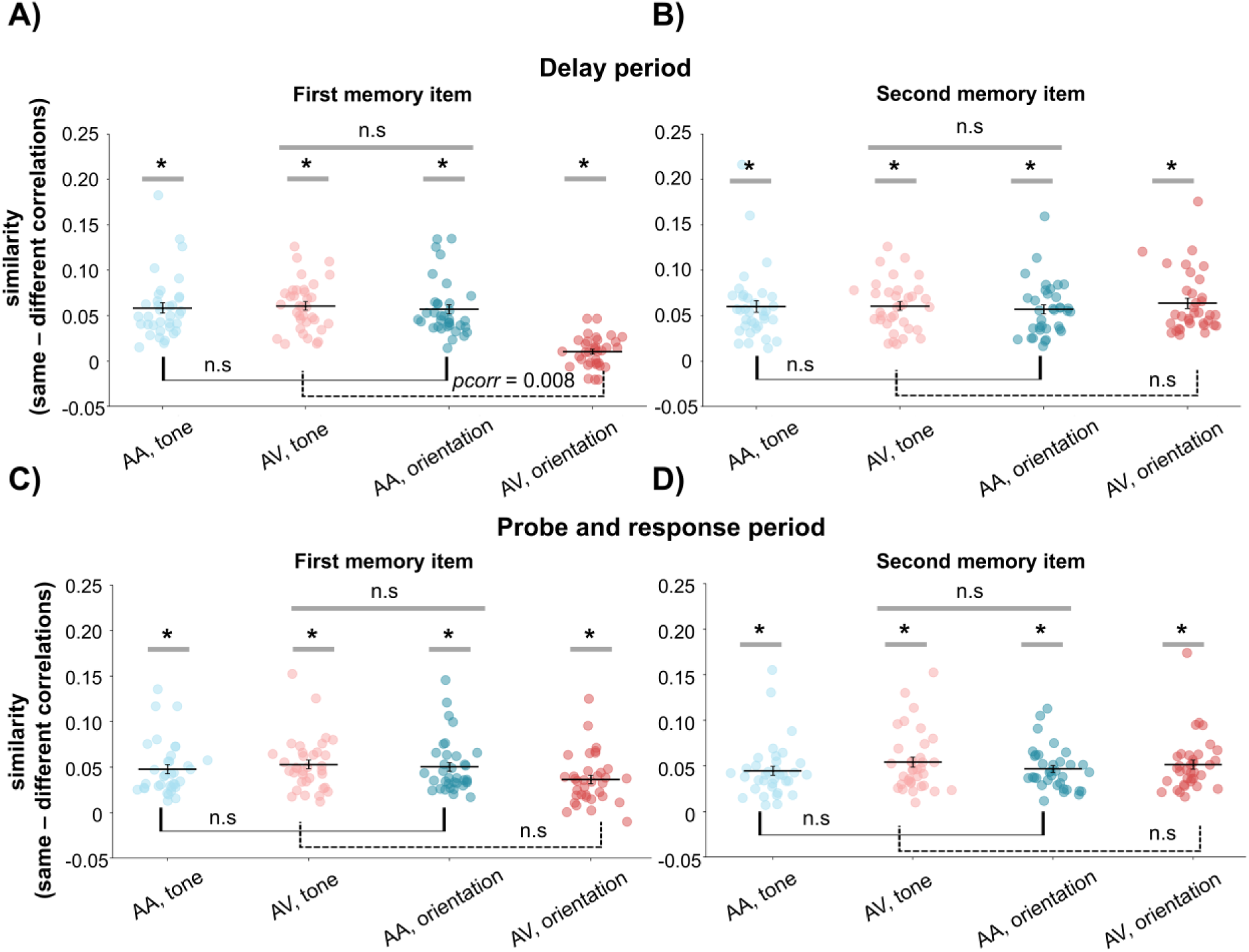
Similarity scores per stimulus feature across conditions. Similarity values are obtained as the difference between “same” vs. “different” correlations for both orientations and tones in attend-auditory (AA) and attend-visual (AV) trials, separately during the delay period; (A) for the first and (B) second item, and during the probe and response period for (C) the first and (D) second item. For statistical tests spearman correlations were Fisher-z-transformed. Circles show individual similarity scores, solid black lines show the average similarity across participants. The error bars show the standard error of the mean. * *p* < 0.05.

Similarly, one-sample t-tests showed that difference scores for tones of the first memory item were significantly different from zero in both attend-auditory (delay: *t*(34) = 9.95, *pcorr* = 0.004, *d* = 1.68; probe-response: *t*(34) = 9.58, *pcorr* = 0.004, *d* = 1.62) and attend-visual (delay: *t*(34) = 13.06, *pcorr* = 0.003, *d* = 2.21; probe-response: *t*(34) = 10.74, *pcorr* = 0.003, *d* = 1.82) trials, indicating that frequency-specific information was present in the neural activity patterns. Compatible results were obtained for tones of the second memory item in both the attend-auditory condition (delay: *t*(34) = 9.1, *pcorr* = 0.002, *d* = 1.54; probe-response: *t*(34) = 8.71, *pcorr* = 0.002, *d* = 1.47) and the attend-visual condition (delay: *t*(34) = 9.27, *pcorr* = 0.001, *d* = 1.57; probe-response: *t*(34) = 9.70, *pcorr* = 0.001, *d* = 1.64). Overall, these results suggest that both orientations and tones of the first and second memory items were represented in working memory throughout maintenance and recall, when attending to a single modality.

Second, we contrasted the difference scores between conditions to test whether task-irrelevant tones (in attend-visual trials) were more strongly represented than task-irrelevant orientations (in attend-auditory trials). However, paired-sample t-tests showed no significant differences in this regard, neither for the features of the first memory item (delay: *t*(34) = -0.55, memory item (delay: *t*(34) = -1.59, *pcorr* = 0.48, *d* = -0.27; probe-response: *t*(34) = -1.14, *pcorr* = 0.78, *d* = -0.11). These results indicate that, although tones and orientations are maintained and, to some degree, reactivated during recall in both attend-conditions, the data do not support the idea that task-irrelevant tones are more strongly represented than task-irrelevant orientations.

Finally, we assessed whether task-relevant features were more strongly represented than task-irrelevant features. In attend-visual trials, difference scores for the first items’ orientations were smaller than those for the tones during the delay period (*t*(34) = -9.84, *pcorr* = 0.008, *d* = -1.66), while they did not differ during the probe-response period (*t*(34) = -2.67, *pcorr* = 0.08, *d* = -0.45). Difference scores for task-relevant orientations of the second item in attend-visual trials were not different from those for the tones, neither during the delay period (*t*(34) = -0.81, *pcorr* = 1, *d* = -0.14) nor in the probe-response period (*t*(34) = -0.42, *pcorr* = 1, *d* = -0.07). In attend-auditory trials, the difference scores for task-relevant tones did not differ from those for orientations, neither in the delay period (first item: *t*(34) = -0.28, *pcorr* = 0.78, *d* = -0.05; second item: *t*(34) = -0.62, *pcorr* = 1, *d* = -0.1), nor the probe-response period (first item: *t*(34) = 0.63, *pcorr* = 1, *d* = 0.1; second item: *t*(34) = 0.45, *pcorr* = 1, *d* = 0.08). These results suggest that the orientations and tones are represented to a similar degree in each selective attention condition. Exceptionally, the tones of the first memory item appeared to be more strongly represented than orientations in the attend-visual trials, yet, this stronger representation disappears during the probe-response period.

## Discussion

Previous research predominantly examined working memory storage of task-irrelevant features within single modalities. Here, we investigated this phenomenon in a multisensory context, employing an audio-visual delayed-match-to-sample paradigm. We provide both behavioral as well as electrophysiological evidence showing that task-irrelevant cross-modal features persist in working memory. Behaviorally, a probe congruency effect indicated that task-irrelevant features impaired recall performance. Corroborating this behavioral effect, representational similarity analyses (RSA) revealed two key results: First, neural activity patterns in selective attention conditions showed greater similarity to the multisensory conjunction condition than to their respective unisensory counterparts, suggesting storage in a conjunction-like format. Second, orientation-*and* tone-specific information was represented in neural activity patterns despite task instructions directing attention to a single modality. These findings expand our understanding of the interplay between multisensory processing and attention in working memory and offer a novel analytical framework to assess neural representations in multisensory contexts.

The observed behavioral probe congruency effect, indicating interference from the involuntary storage of task-irrelevant features, replicates previous findings extensively reported in unisensory contexts (Ecker et al., 2013; Fischer et al., 2024.; Gu et al., 2020; Joseph et al., 2015; Logie et al., 2011) as well as recent evidence from a multisensory working memory paradigm (Arslan et al., 2025). In particular, for temporally and spatially aligned audio-visual features, the encoding of task-irrelevant features was expected, given that these bottom-up factors have been shown to promote cross-modal feature integration (Stein & Stanford, 2008; Van Der Burg et al., 2008, 2010; Wallace et al., 2004). However, considering previous reports of attenuated multisensory integration when features occur in separate spatial locations (Gondan et al., 2005; Teder-Sälejärvi et al., 2005), we expected reduced interference from irrelevant features in spatially disparate trials. In contrast, the probe congruency effect was even stronger in the attend-visual condition when tones were presented laterally, while visual targets remained centrally located. The disruptive influence of lateralized tones was also observed in a decreased overall accuracy of attend-visual trials in spatially disparate compared to spatially compatible trials. Notably, the lateralization of the tones did not affect performance in attend-auditory trials. We propose that lateralized sounds in spatially disparate attend-visual trials, varying unpredictably in their spatial location from trial to trial, may have captured exogenous spatial attention (Blauert, 1996; Spence et al., 1998; Spence & Driver, 1997). This attentional capture would have diverted resources away from the centrally presented visual stimuli, thereby explaining both the overall reduced accuracy and the increased probe congruency effects observed in attend-visual trials under spatially disparate conditions.

Consistently, even sounds spatially segregated from visual targets have been shown to elicit an Auditory-Evoked Contralateral Occipital Positivity (ACOP) - an ERP component associated with the orienting of visual attention to a sound’s location - though to a lesser extent than spatially aligned sounds (Hillyard et al., 2016). In addition, visual attention has been shown to spread to task-irrelevant sounds, even when presented at different locations, as reflected in an enhancement of auditory cortex activity (Busse et al., 2005). Notably, the feature-level RSA revealed an intriguing, though exploratory, finding that further corroborates this line of argumentation: that is, task-irrelevant tone frequencies of the first item were represented more strongly than task-relevant visual orientations during the delay period of attend-visual trials (Figure 8A, AV tones vs. AV orientations). One tentative explanation for this effect could be that the spatial location of the first sound could not be anticipated, as sound locations varied unpredictably from trial to trial. Consequently, the first sound may have elicited stronger exogenous attentional capture, leading to a more robust initial encoding of the auditory feature. For the second item, which retained the same spatial configurations as the first item, no such heightened representation of task-irrelevant sounds emerged, suggesting that attentional capture diminished once the spatial location of the sound was known. Altogether, these findings support the notion that task-irrelevant auditory stimuli can affect the processing of visual stimuli, even in the absence of spatial congruency.

Thus, our findings challenge the commonly held assumption that spatial segregation attenuates multisensory interactions. While a substantial body of behavioral (Delong & Noppeney, 2021; Frassinetti et al., 2002; Spence et al., 1998; Zimmer & Macaluso, 2007) and neuroscience (Gondan et al., 2005; Teder-Sälejärvi et al., 2005) literature shows that spatial congruency enhances crossmodal interactions, such effects – in particular on the behavioral level – typically arise when location is task-informative or partially task-relevant (reviewed by Spence, 2013). In such cases, spatially coinciding task-irrelevant auditory, visual, or tactile stimuli often function as spatial cues, improving target detection or discrimination performance (Goldring et al., 1996; Hughes et al., 1994, 1998; Sambo & Forster, 2009; Spence et al., 1998), especially when participants must sample information from different possible target positions. In contrast, spatial segregation between audio-visual features failed to exert an effect on behavioral performance in studies focusing on stimulus identification or temporal perception (Doyle & Snowden, 2001; Keetels & Vroomen, 2005). Critically, in the present study, we deliberately rendered the spatial information task-irrelevant, following the rationale that demonstrating perceptual binding effects – rather than decision-level integration – requires the dimension on which subjects make a report to differ from the dimension that is intended to create the binding (Bizley et al., 2016). This design choice may have diminished the effects of spatial congruency on multisensory interactions.

The behavioral interference from task-irrelevant features is further corroborated by our neural findings. First, by comparing neural activity pattern similarities across conditions, we found that both attend-auditory and attend-visual trials were initially more similar to the conjunction condition than to their corresponding unimodal conditions (i.e., auditory-only and visual-only, cf. Figure 7). This suggests that the representational structure in these selective attention conditions reflects integrated feature conjunctions rather than individual features. While higher similarity to the conjunction condition during stimulus presentation is not surprising — given the physical similarity of stimuli during those stages — we focus our interpretation on the delay and response period following probe onset. This revealed distinct representational dynamics of attend-auditory and attend-visual trials across time, showing that task-irrelevant visual and auditory features were not processed equivalently in the present study: In the attend-auditory condition, representations initially resembled integrated conjunctions, but this similarity diminished later in the delay. Crucially, higher similarity to the auditory-only condition never emerged, suggesting only partial filtering of irrelevant orientations – this aligns with the behavioral evidence of persistent interference at recall. In contrast, in the attend-visual condition, similarity to the conjunction condition remained high throughout the delay period, indicating that task-irrelevant tones were not effectively filtered out. Notably, this effect was stronger in spatially disparate trials, where tones were lateralized, mirroring the behavioral findings that lateralized, task-irrelevant tones were more disruptive than centrally presented ones.

At recall, the attend-auditory condition again showed equal similarity to both the conjunction and auditory-only conditions, in line with the interpretation that orientation information was partially filtered out and participants remained focused on auditory features. In contrast, after probe-offset, attend-visual trials exhibited stronger similarity to the conjunction condition than to visual-only trials. This pattern may reflect two non-exclusive mechanisms: either participants failed to suppress probe tones effectively, thus, encoding them into working memory, or memorized tone features were reactivated during recall. To further investigate these possibilities, we conducted an exploratory follow-up analysis to assess whether activity patterns contain orientation- or tone-specific information. Thus, while the primary analysis assessed representational similarity on the condition level, the feature-level RSA resembles a decoding approach, providing direct neural evidence of orientation- or tone-specific information.

In fact, feature-level RSA showed that memory representations contained both orientation- and tone-specific information throughout the delay period and recall, irrespective of the attended modality. This pattern alludes to findings from the episodic memory literature, reporting the incidental reactivation of task-irrelevant cross-modal features encoded during stimulus presentation (Nyberg et al., 2000). The latter is in line with the proposal that retrieval of one component results in the reactivation of other components present at encoding (Alvarez & Squire, 1994). However, given that task-irrelevant tones were not more strongly represented than task-irrelevant orientations (apart from a transient effect earlier in the delay for the first item), we cannot exclude that the greater similarity of attend-visual to conjunction trials during recall (which is not present for attend-auditory trials, see recall period in Fig 6C and D vs. 6A and B) is primarily due to attentional processing of the probe tones rather than a reactivation of memorized auditory features.

Further, we did not observe a general increase in representational strength for task-relevant over task-irrelevant features (cf. Figure 8). This contrasts with prior studies of unisensory working memory, which typically show enhanced representation of task-relevant information (Chen et al., 2021) or fail to decode task-irrelevant feature altogether (Bocincova & Johnson, 2019; Yu & Shim, 2017). Critically, rather than representing a limitation, this discrepancy may reflect inherent differences in representational dynamics associated with multisensory processing. Specifically, multisensory integration is typically characterized by nonlinear neural responses, which differ significantly from the sum of responses to unisensory inputs (Fort, 2002; Foxe et al., 2000; Giard & Peronnet, 1999; Murray et al., 2005; Senkowski et al., 2007, 2011; Talsma & Woldorff, 2005). Such nonlinearities support the view that qualitatively distinct neural mechanisms for multisensory stimuli may promote stronger feature binding - even under conditions of modality-specific attention (Talsma & Woldorff, 2005) - thereby facilitating the persistent representation of task-relevant *and* task-irrelevant features observed here. However, we should note that we did not observe significant deviations from additivity during the encoding period of the first item in a supplementary analysis of univariate ERP responses (see section 1.3 in supplementary material). This null result may reflect differences in task demands and cognitive state, given that the present study involved encoding into working memory rather than simple detection, as employed in most studies adopting additive models. Importantly, this does not preclude that multisensory binding took place in the present study. Indeed, others have similarly reported that audiovisual ERPs follow an additive principle, while supra-additive interactions only emerged in multivariate activity patterns (Buhmann et al., 2024). These findings highlight the limitations of classical additive models to reveal binding effects in more complex tasks, such as working memory paradigms, and underscore the values of multivariate approaches like RSA.

Future studies should explicitly assess the neural dynamics underlying cross-modal as opposed to unisensory feature bindings in working memory. To date, behavioral evidence points to a multisensory benefit for memory (Lehmann & Murray, 2005; Matusz et al., 2015; Murray et al., 2005; Thelen et al., 2012, 2015; Y. Xie et al., 2017, 2021; Y. J. Xie et al., 2019) which may yield performance advantages over unisensory bindings under certain conditions (but see also Allen et al., 2009; Guazzo et al., 2020). This aligns with the notion that multisensory information can give rise to representations that are richer in physical features (Nairne, 1990; Neath, 2000), even when attention is selectively directed to a single modality (Lehmann & Murray, 2005).

A more trivial explanation for the sustained storage of task-irrelevant features could be that participants never tried to selectively attend to only one modality. However, multiple findings argue against this interpretation: First, as shown in our previous work (Arslan et al., 2025), the probe congruency effect was significantly larger in the conjunction condition compared to the selective attention conditions (see section 2 in supplementary material) – an outcome that would not be expected if both modalities were consistently attended. Second, condition-level RSA results, at least for the attend-auditory condition, indicate partial filtering of task-irrelevant visual features. Given that attend-auditory and attend-visual trials were randomly inter-leaved within one block at a high pace, it seems unlikely that participants strategically alternated between dual- and single-modality attention. Rather, the lack of or attenuated filtering in attend-visual trials is more plausibly explained by the involuntary spread of attention to lateralized sounds (Busse et al., 2005; Hillyard et al., 2016). This is further corroborated by participants answers in a follow-up questionnaire, indicating that they adhered to the selective attention instructions in both conditions. On a different note, one could argue that the filtering of task-irrelevant orientations is caused by participants looking away in attend-auditory trials. Although the present study did not include eye-tracking, 30 out of 35 participants indicated in a follow-up questionnaire that they maintained their gaze in the center of the screen in attend-auditory trials. Importantly, a control analysis, excluding those five participants who noted that they sometimes failed to fixate on the center of the screen, replicated the original pattern of results (see section 4 in supplementary material).

Finally, returning to the discrepancy with prior unisensory decoding studies, we note that the inability to decode task-irrelevant features in those studies (Serences et al., 2009; Yu & Shim, 2017) does not necessarily imply their complete absence from working memory. For instance, recent work has shown that uncued – i.e., task-irrelevant – color information can be recovered from impulse-evoked EEG activity (Kandemir et al., 2024), demonstrating that non-prioritized information may not be completely lost. Notably, decoding studies that assess the neural fate of task-irrelevant features typically lack corresponding behavioral measures and analyses of incidental feature reactivation during recall. Combing neuroimaging, impulse-perturbation paradigms (Wolff et al., 2015, 2020) and manipulations of irrelevant-change effects (e.g., Fischer et al., 2024; Joseph et al., 2015), future studies could directly address the link between neural representations and behavioral interference.

### Limitations

While our findings offer both methodological and theoretical advances to the study of multisensory working memory, some limitations should be noted. First, the relatively limited number of trials per condition and consequentially per feature value (i.e., individual orientations and tone frequencies) restricted our analyses to coarse distinction at the feature level (i.e., left versus right orientations and low versus high frequencies). This constrains the granularity of our interpretations. Second, the present results do not provide any insights into the spatial localization of effects. Thus, we cannot make any claims as to where in the brain incidentally formed multisensory representations reside – a question that future studies using complementary neuroimaging methods such as fMRI might address.

## Conclusion

Our behavioral and RSA results demonstrate that audiovisual working memory representations inherently include task-irrelevant features, highlighting the automaticity of cross-modal binding, despite the attentional prioritization of one modality over the other. Critically, we found behavioral and neural indices of task-irrelevant feature storage to be robust, even when audiovisual features were segregated at encoding. This pattern implies that bottom-up driven integration at encoding may provide an effective scaffold for persistent multisensory representations, which may be qualitatively distinct from those formed in unisensory contexts – though future work is needed to explicitly clarify the nature of potential differences in unisensory and multisensory bindings.

We also observed differential representational dynamics for task-irrelevant auditory versus visual features, likely related to attentional capture by lateralized auditory stimuli. This points to a role for exogenous attention in modulating how multisensory features are bound and maintained in working memory.

Taken together, the data support a hierarchical model in which modality-specific prioritization typically occurs during maintenance, while cross-modal links remain intact and can be dynamically retrieved - either incidentally or in line with task demands. This interpretation aligns with our previous work (Arslan et al., 2025), which showed modality-specific prioritization during the delay period and greater integrative processing during recall. Together, our results point to a flexible architecture for multisensory working memory, in which selective attention interacts with object representations (Ernst et al., 2013; O’Craven et al., 1999) that include both task-relevant and task-irrelevant features.

### Data and Code Availability

All behavioral and processed EEG data as well as analysis scripts and code supporting the results are provided at this link: https://osf.io/xakqw/. Raw EEG data will be shared upon manuscript acceptance.

### Declaration of Competing Interests

The authors have no competing interests to declare.

### CRediT statement

Conceptualization: LIK (lead), CA, DS, and SG (support); Methodology: LIK; Formal analysis: CA (lead), LIK (support); Writing – Original Draft: CA (lead), LIK (support); Writing – Review & Editing: LIK and CA (lead), DS, EW and SG (support); Visualization: CA (lead), LIK (support); Investigation: CA; data curation: CA; Supervision: LIK; Project administration: LIK (lead), CA (support); Resources: EW.

## Acknowledgements

The authors would like to thank Tobias Blanke for programming the experiment and Mareike Vienken, Clara Weitzenböck, Rama Alnabulsy, Hanna Margarita Torkler and Jane Westedt for their support in collecting the data. In addition, the authors are grateful to Eren Günseli for the suggestion to conduct a feature-level RSA and for helpful comments on how to implement it.

## Notes

https://osf.io/xakqw/

### Competing Interest Statement

The authors have declared no competing interest.

