## Supplementary material for "Tracking neural representations of attended and unattended features in multisensory working memory over time"

**Author Note**

Ceren Arslan,
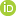
<https://orcid.org/0000-0001-6905-1832> https://orcid.org/0000-0003-0601-0747

Daniel Schneider,
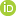
<https://orcid.org/0000-0001-6905-1832> https://orcid.org/0000-0002-2867-2613

Stephan Getzmann,
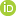
<https://orcid.org/0000-0001-6905-1832> https://orcid.org/0000-0002-6382-0183

Edmund Wascher,
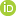
<https://orcid.org/0000-0001-6905-1832> https://orcid.org/0000-0003-3616-9767

Laura-Isabelle Klatt,
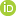
<https://orcid.org/0000-0001-6905-1832> https://orcid.org/0000-0002-5682-5824

**Correspondence**

Ceren Arslan, Leibniz Research Centre for Working Environment and Human Factors, Ardeystraße 67, 44139 Dortmund, Germany.

**Supplementary Materials**

1. **Assessment of multisensory integration processes using additive models**

To investigate crossmodal interactions, we used an additive model framework (Besle et al., 2004; Cappe et al., 2010; Fort, 2002; Giard & Peronnet, 1999; Senkowski et al., 2011; Stekelenburg & Vroomen, 2007; Talsma & Woldorff, 2005), contrasting the conjunction condition ERP with the summed responses of the unisensory conditions. The rational of this additive model implies that if the auditory (A) and visual (V) features of an audiovisual (AV) compound stimulus are processed entirely independent from each other, the neural response to AV should be equal to the sum of A and V. On the contrary, any supra-additive or sub-additive effect should be attributed to interactions between the two modalities (Besle et al., 2004; Cappe et al., 2010; Fort, 2002; Giard & Peronnet, 1999; Senkowski et al., 2011; Stekelenburg & Vroomen, 2007; Talsma & Woldorff, 2005). However, additive models can result in spurious crossmodal interactions unless certain methodological caveats are met (Besle et al., 2004; Giard & Besle, 2010):

First, to avoid a continuous deactivation of a given sensory cortex, resulting in an artificial increase of crossmodal effects, we randomly presented ‘auditory-only’, ‘visual-only’ and ‘conjunction’ trials in A-Blocks. In contrast, attend-auditory and attend-visual trials, which necessarily required the instruction to direct attention to one modality, were grouped and randomly presented in B-Blocks. This ensured that all trial types included in the additive model (A-Blocks) were randomly and equiprobably delivered across all modalities (Besle et al., 2004).

Second, to eliminate anticipatory effects common to all conditions, the interval between the trial-onset cue and the first stimulus was randomly jittered. In addition, we included aborted trials, in which participants expected a regular trial to occur but in which no stimulus appears (Busse & Woldorff, 2003, Senkowski et al., 2011). Specifically, in 30% of trials in A-Blocks, the variable inter-stimulus-interval after the onset-cue was followed by a white fixation cross, which had the same duration as a memory item (500 ms). At this rate, no-stimulus events have been shown to not elicit a physiological omission response (Busse & Woldorff, 2003). Considering that these events might have been an opportunity to ‘rest’ between actual task trials, we also included these events in B-Blocks.

By applying this model, we aimed to examine how spatial manipulation influences multisensory interactions. Specifically, we hypothesized that: (1) if spatial compatibility between auditory and visual features enhances multisensory interactions, this should manifest as a supra-additive or sub-additive effect; and (2) if spatial segregation successfully disrupts feature binding, then this supra- or sub-additive effect should be reduced in spatially disparate trials.

**1.1 Preprocessing**

Preprocessing followed the same procedure as described in the main manuscript, with specific adjustments for the ERP time window: Considering that only a short time window can be considered (see also 1.2), the data was epoched between -500 to 500 ms, relative to the onset of the first item. On average, 4.74 channels were rejected (*SD* = 1.46, range = 1 to 7), and 27 components (*SD* = 5.98, range = 13 to 37) were excluded. On average, 141.66 trials remained in conjunction (*SD* = 7, 98%), 141.54 trials in auditory-only (*SD* = 6.46, 98%), 141.37 trials in visual-only (*SD* = 7, 98%) conditions, and 188.29 no-stimulus events (*SD* = 9, 98%) remained in pre-processed data.

**1.2 ERP analyses**

First, to eradicate the effect of anticipatory activity (Senkowski et al., 2007; Talsma & Woldorff, 2005; Teder-Sälejärvi et al., 2002), the grand-average ERP of aborted trials (i.e., no-stimulus events) was subtracted from the ERPs of each condition (i.e., conjunction, auditory-only, visual-only) To explore whether spatial congruency modulated crossmodal interactions, the sum of auditory- and visual-only waveforms was subtracted from the conjunction condition separately for spatially compatible and disparate trials:

$$eq \left( 1 \right): {(ERP conjunction}_{compatible})-({ERP Aonly}_{compatible}+{ERP Vonly}_{compatible} )$$

$$eq \left( 2 \right): {(ERP conjunction}_{disparate})-({ERP Aonly}_{disparate}+{ERP Vonly}_{disparate} )$$

If the spatial disparity between audio-visual features attenuated crossmodal interactions, the amplitude difference in the spatially compatible condition is greater than the difference in the spatially disparate condition.

Mean amplitudes for each condition were calculated using a cluster of central electrodes, including C1/C2/Cz, CP1/CP2/CPz, FC1/FC2, and posterior electrodes, including PO3/PO4, P3/P4/Pz, and O1/O2/Oz (Senkowski et al., 2007). Differences in mean amplitudes were submitted to one-sample t-tests separately for spatially compatible and disparate conditions and for central and parietal electrode clusters. The statistical analysis focused on the time interval between 0 to 200 ms, considering that common activity, such as motor processes, response selection, or semantic processes, usually emerges 200 ms post-stimulus. The latter would effectively be subtracted twice in the [AV - (A + V)] model, potentially resulting in spurious cross-modal interactions (Besle et al., 2004; Fort, 2002).

**1.3 Results**

ERP amplitudes in the conjunction condition did not significantly different from the sum of the auditory-only and visual-only conditions, neither in spatially compatible (central cluster: *t*(34) = -0.54, *pcorr* = 1, *d* = -0.1; posterior cluster: *t*(34) = -1.65, *pcorr* = 0.42, *d* = -0.28) nor in spatially disparate trials (central cluster: *t*(34) = -1.59, *pcorr* = 0.37, *d* = -0.27; posterior cluster: *t*(34) = -0.03, *pcorr* = 0.98, *d* = -0.01) (see supplementary figure 1). Given that no supra- or sub-additive effects were established within each condition, no further analyses comparing ERP differences between spatially compatible and spatially segregated trials were conducted.

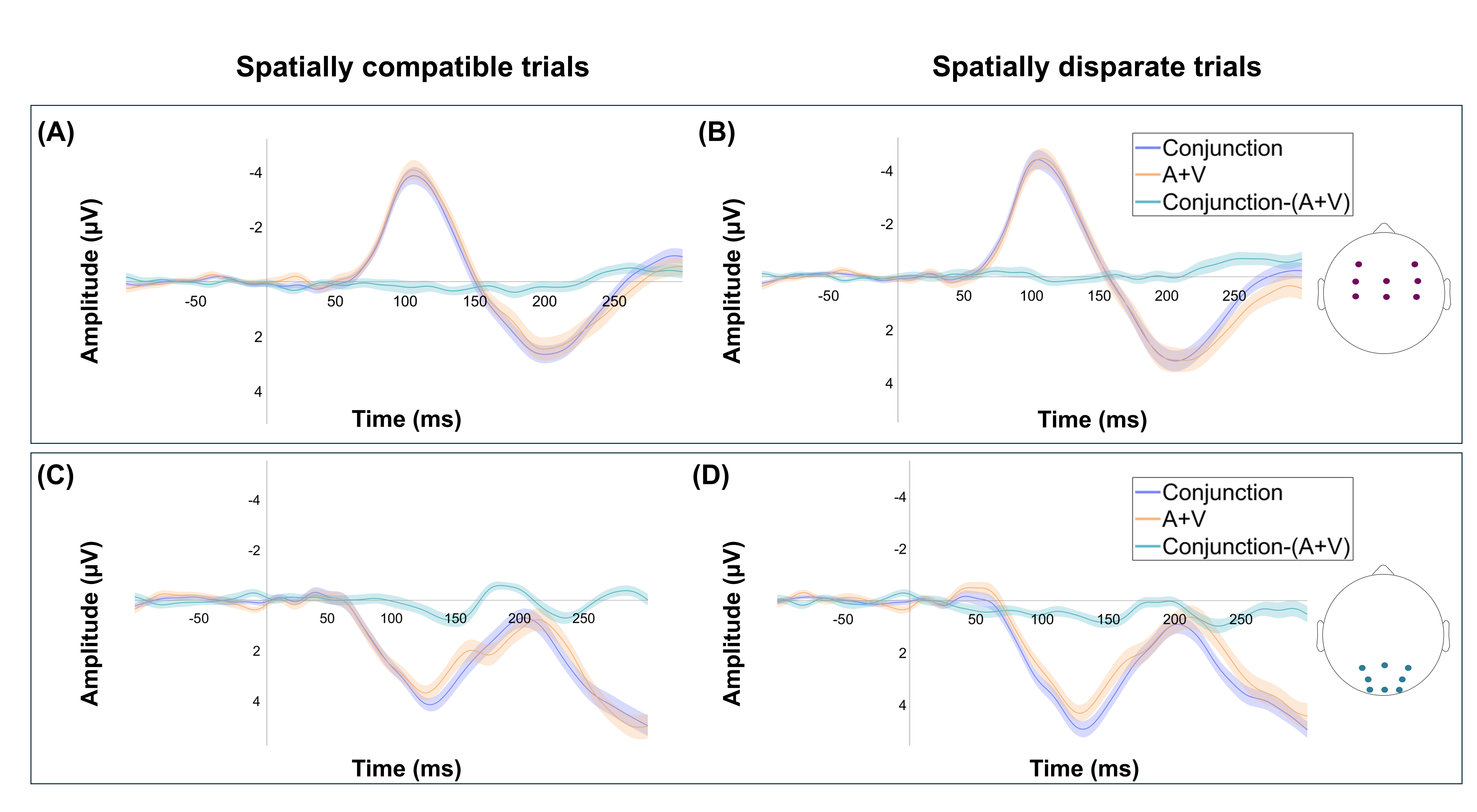

**Supplementary Figure 1.** Grand-averaged ERP waveforms, illustrating additive model contrasts. Panels (A) and (C) illustrate ERP waveforms for spatially compatible trials, while panels (B) and (D) illustrate ERP waveforms for spatially disparate trials for central (top row) and posterior (bottom row) electrode clusters, respectively.

2. **Probe congruency effects under unimodal vs bimodal attention**

In our previous work, we showed that congruency effects were significantly stronger when attending to both (i.e., conjunction condition) rather than to only one modality (Arslan et al., 2025). This is consistent with the assumption that when both modalities are attended, partial changes of the probe features in only one modality are more detrimental to performance than when participants selectively attend to either audition or vision. The present supplementary analysis examines analogous differences in the size of the congruency effect between conditions.

Using paired-sample t-tests (see supplementary table 1), we contrasted the differences in accuracy between congruent and incongruent trials between the conjunction condition and the two selective attention conditions. Note that only ‘no’ response trials were considered in this analysis to account for the unequal distribution of “yes” trials across congruent and incongruent probes in the conjunction and the selective attention conditions (for details, see Figure 2 in the main manuscript).

Probe congruency effects in accuracy were significantly greater in the conjunction condition compared to the two selective attention conditions (*t*(34) = 3.37, *p* = 0.002, *d* = 0.57).

When contrasting the conjunction condition separately with the attend-auditory and the attend-visual condition, respectively, the results strongly mirror the condition-level RSA results. That is, the probe congruency effect was significantly diminished in attend-auditory trials (*M* = 5.79, *SD* = 8.66) compared to conjunction trials (*M* = 15.87, *SD* = 15.28), (*t*(34) = 3.81, *p* < 0.01, *d* = 0.64), yet, probe congruency effects did not significantly differ between the attend-visual (*M* = 5.50, *SD* = 9.54) and the conjunction trials (*M* = 7.30, *SD* = 12.37), (*t*(34) = 0.83, *p* = 0.42, *d* = 0.14) (for probe types chosen for each contrast see Supplementary Table 1). This suggests that participants were less able to suppress the task-irrelevant auditory features. Critically, this was not due to participants strategically attending to both modalities in attend-visual trials (see follow-up questionnaire ratings below).

For congruency effects in response times, no significant differences emerged between conditions (all *p* > 0.85).

**Supplementary Table 1.** Probe types included in the calculation of probe congruency effects.

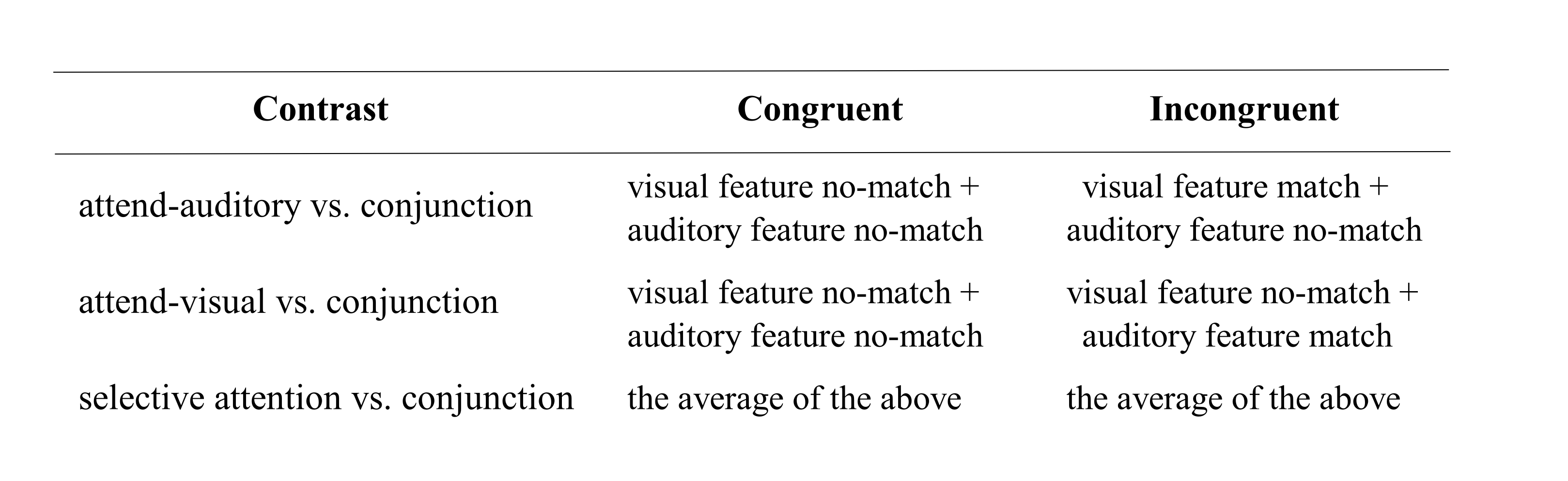

3. **Follow-up questionnaire**

To assess if the participants adhered to the task instructions that required them to attend only to one modality in the selective attention conditions, we asked the following questions in a follow-up questionnaire: 1) "In the auditory task, were you able to focus only on the sounds, and did you try to ignore the visual stimuli?", 2) "In the visual task, were you able to focus only on the visual stimuli, and did you try to ignore the sounds?". Participants reported their answers on a scale from 1 (never) to 5 (always). The self-reported answers indicated that participants adhered to the selective attention instructions and attempted to attend only to the task-relevant modality while ignoring the task-irrelevant features from the other modality. Supplementary Table 2 demonstrates the means and standard deviations of participants’ ratings to those questions. Additionally, the participants were asked a ‘yes or no’ question 3) ‘Did you continue to look at the center of the screen in the auditory conditions?’ with the possibility of reporting a more detailed answer. Overall, five participants answered ‘no’, including one reporting, ‘I tried, but sometimes I looked in the corner.’ and one writing, ‘In the beginning, I did it very well, but in the end, I did not focus on the fixation cross.’

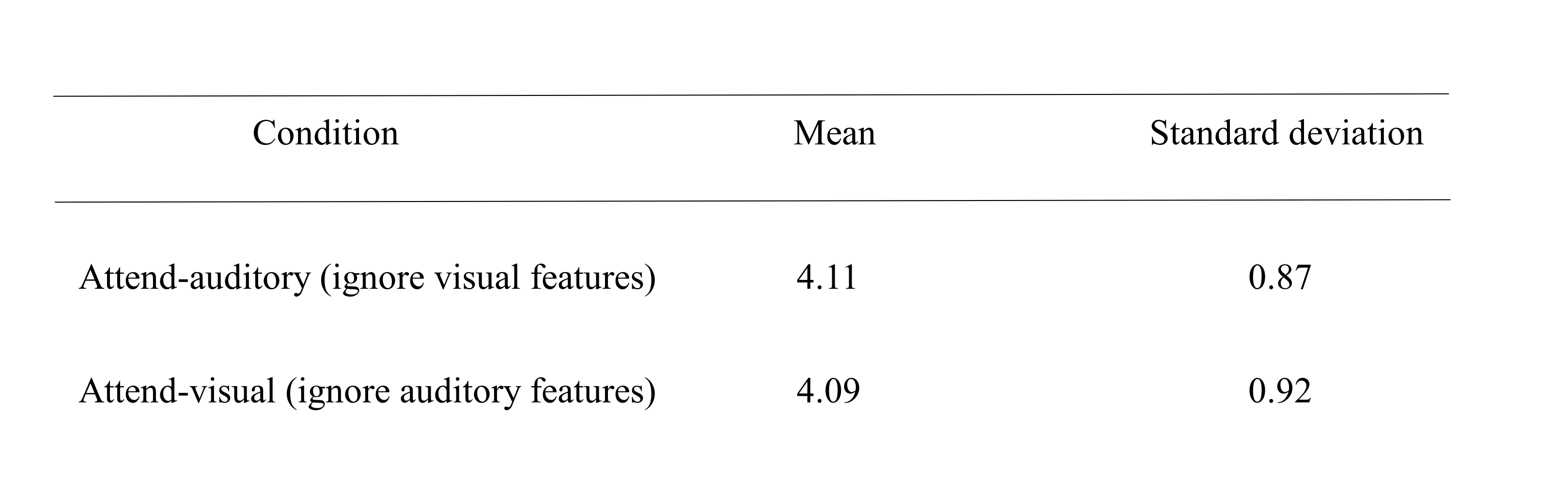
**Supplementary Table 2.** Follow-up questionnaire ratings.

Note: Participants indicated to what extend they were able to focus on the task-relevant modality and attempted to ignore the task-irrelevant modality on a scale from 1 (never) to 5 (always).

4. **Representational similarity between task conditions**

Considering that five participants reported that they did not fixate on the center of the screen during the auditory-only and attend-auditory trials, condition-level RSA analyses were re-run without those participants. The goal of these supplementary analyses was to compare the selective conjunction and selective unimodal contrasts. The same procedure described in the manuscript was followed for these tests (see 2.6.2.1 Representational similarity between task conditions). Results replicate the pattern observed with the full sample as reported in the manuscript (see 3.2 Representational similarity between conditions), with slight differences in the time range of the significant clusters

For spatially compatible trials, RSA revealed significantly higher pattern similarity of attend-auditory trials with the conjunction than with the auditory-only condition. Significant clusters (*p* < .05) emerged during encoding as well as parts of the delay interval (i.e., 80 to 1150 ms, between 1910 to 2070 ms, and between 2470 to 2690 ms, (see supplementary figure 2A). For spatially disparate feature presentation, a similar pattern emerged with significant clusters ranging from 80 to 1220 ms, 1760 to 2200 ms, and 2460 to 2780 ms (*p* < .05) (see supplementary figure 2B).

For spatially compatible attend-visual trials, RSA revealed higher similarity to the conjunction condition than to the visual-only condition. The effect was sustained throughout the majority of the trial with significant clusters ranging from 40 to 2830 ms and between 3530 to 4220 ms (*p* < .05) (see supplementary figure 2C). A compatible pattern was obtained for spatially disparate trials with significant clusters ranging from 45 to 2890 ms and from 3530 to 4290 ms (*p* < .05) (see supplementary figure 2D).

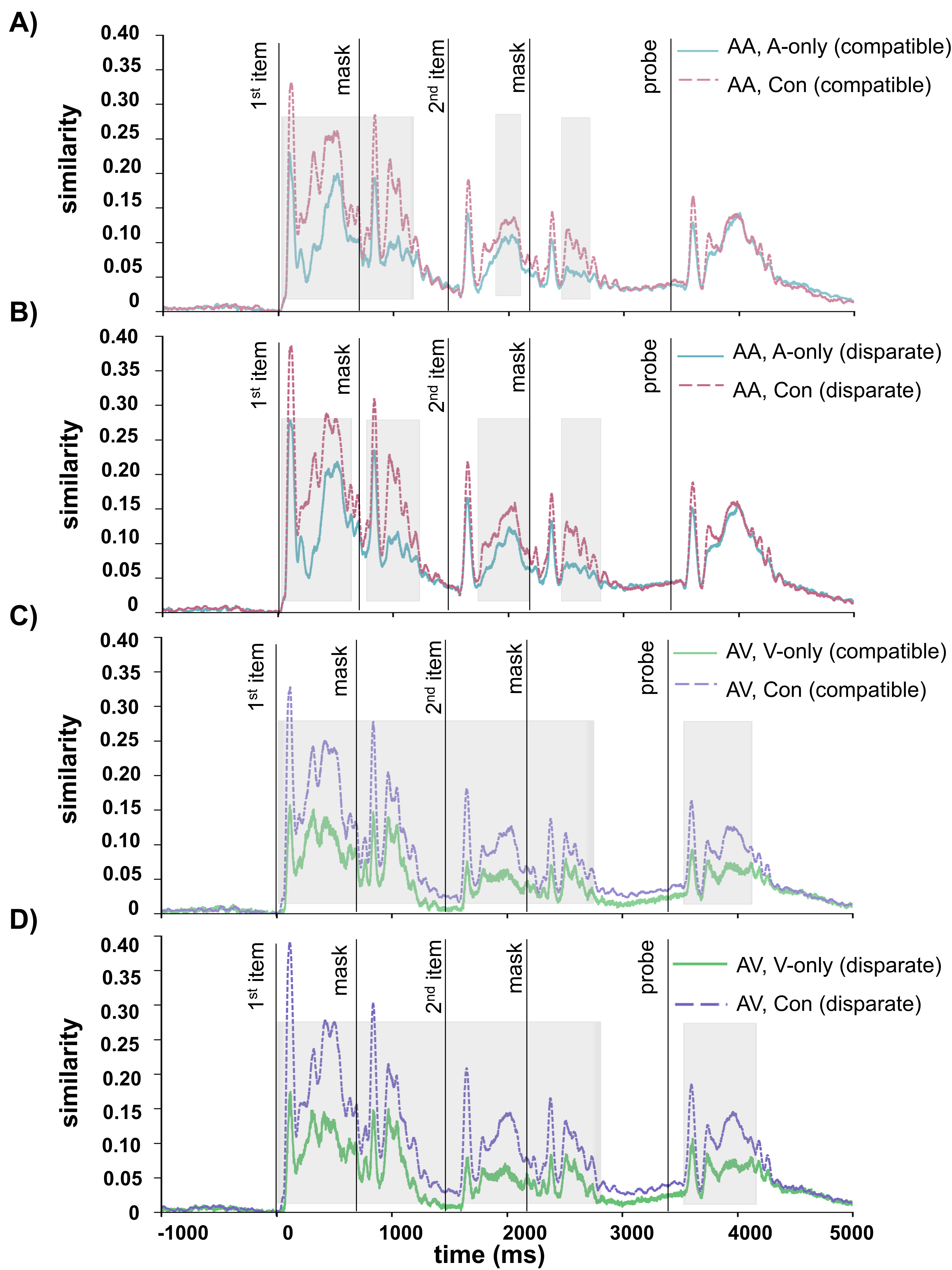

**Supplementary Figure 2.** Similarities of neural activity patterns between task conditions for spatially compatible and disparate trials. Panels A and B show Spearman correlations between attend-auditory (AA) and auditory-only (A-only) trials (blue solid line) and between attend-auditory (AA) and conjunction (Con) trials (pink dashed line) for spatially compatible and for spatially disparate conditions, respectively. Panels C and D show between attend-visual (AV) and visual-only (V-only) trials (green solid line) and between attend-visual (AV) and conjunction (Con) trials (purple dashed line) for (C) spatially compatible and (D) for spatially disparate trials. Shaded areas show the time periods in which the pairwise comparisons significantly differed.
